# Kinetic and thermodynamic analysis defines roles for two metal ions in DNA polymerase specificity and catalysis

**DOI:** 10.1101/2020.10.20.347138

**Authors:** Shanzhong Gong, Serdal Kirmizialtin, Adrienne Chang, Joshua E. Mayfield, Yan Jessie Zhang, Kenneth A. Johnson

**Affiliations:** Department of Molecular Biosciences, The University of Texas at Austin, 2500 Speedway, MBB 3.122, Austin, Texas 78712; Chemistry Program, Science Division, New York University Abu Dhabi, Abu Dhabi, United Arab Emirates

**Keywords:** DNA replication, magnesium, molecular dynamics, conformational change, enzyme catalysis, transient kinetics, induced-fit, enzyme specificity, metal ions

## Abstract

We examined the roles of Mg^2+^ ions in DNA polymerization by kinetic analysis of single nucleotide incorporation catalyzed by HIV reverse transcriptase and by molecular dynamics simulation of Mg^2+^ binding. Binding of the Mg-nucleotide complex induces a conformational change of the enzyme from open to closed states in a process that is independent of free Mg^2+^ concentration. Subsequently, the second Mg^2+^ binds weakly to the closed state of the enzyme-DNA-Mg.dNTP complex with an apparent *K*_*d*_ = 3.7 mM and facilitates the catalytic reaction. This weak binding of the catalytic Mg^2+^ is important to maintain fidelity in that the Mg^2+^ samples the correctly aligned substrate without perturbing the equilibrium at physiological Mg^2+^ concentrations. The binding of the catalytic Mg^2+^ increases nucleotide specificity (*k*_*cat*_/*K*_*m*_) by increasing the rate of the chemistry and decreasing the rate of enzyme opening allowing nucleotide release. Changing the free Mg^2+^ concentration from 0.25 to 10 mM increased nucleotide specificity (*k*_*cat*_/*K*_*m*_) by 12-fold. Mg^2+^ binds very weakly to the open state of the enzyme in the absence of nucleotide (*K*_*d*_ ≈ 34 mM) and competes with Mg.dNTP. Analysis based on publish crystal structures showed that HIV RT binds only two metal ions during incorporation of a correct base-pair. MD simulations support the kinetic studies suggesting weak binding of the catalytic Mg^2+^ in open and closed states. They also support the two-metal ion mechanism, although the polymerase may bind a third metal ion in the presence of a mismatched nucleotide.

## Introduction

Metal ions play critical roles in many biological activities including DNA replication, DNA repair, and transcription as well as other phosphoryl-group transfer reactions. They stabilize the structures of proteins and nucleic acids, and promote the catalytic activities (1). Magnesium ion (Mg^2+^) serves as the primary metal ion for catalysis, due to its natural abundance *in vivo* and restricted coordination geometry conferring high stereoselectivity (2). The role of metal ions in DNA polymerization and hydrolysis was first described by Steitz in 1993 (3) who proposed a two-metal-ion mechanism in which one metal ion forms a tight complex with the incoming nucleotide by coordinating with non-bridging oxygens from all three phosphates (4,5). A second metal ion reduces the pK_a_ of the 3’-OH group for polymerization (or of a water molecule for hydrolysis), thereby activating the nucleophile and bringing it close to the α-phosphate at the reaction center. The coordinated action of two metal ions, water molecules, and several surrounding acidic residues helps to stabilize the transition state (1,6). After polymerization, Mg-pyrophosphate (Mg-PPi) is released from the enzyme (3,7). The two-metal-ion mechanism is supported by many crystal structures of DNA polymerases (8). However, crystal structures only provide a static picture of the active site, do not reveal weakly bound metal ions or their dynamic movements, and do not reveal the pathway or thermodynamics of the reaction. Moreover, dideoxy-terminated primer, calcium ions, or non-hydrolysable nucleotide analogs are usually used in crystal structures to prevent catalysis and these may disrupt the active site geometry and the conformational state of the enzyme. Recently it has been proposed that a third metal ion may be required for catalysis (9,10). The third metal ion is seen predominantly in DNA repair enzymes and may be associated with stabilizing the product pyrophosphate (PPi) (11) but the role of a possible third metal ion in catalysis remains unresolved (12) and has not been seen in higher fidelity enzymes.

Enzyme mechanism and specificity are kinetic phenomena that cannot be addressed by structural studies alone. Rather, structural studies provide the framework for kinetic and mechanistic analysis, so the two approaches together provide new insights. To further understand the role of Mg^2+^ in catalysis and specificity, studies on the dynamics of the metal ions under biologically relevant conditions are required. HIV reverse transcriptase (HIVRT) belongs to the A family of moderate-to high-fidelity enzymes. It serves as a good candidate for studying the two-metal-ion mechanism because kinetic characterization of single nucleotide incorporation has established the mechanistic basis for polymerase fidelity and have defined the role of a nucleotide-induced conformational change step in specificity (13)(14). These studies have established the following minimal pathway for nucleotide incorporation for HIVRT.

Specificity for cognate nucleotide incorporation by HIVRT is a function on an induced-fit mechanism where *k*_*cat*_/*K*_*m*_ is defined by the rate of the fast conformational change to the closed state (*k_2_*) divided by the *K*_*d*_ for the weak binding of nucleotide to the open state of the enzyme (15,16) as shown in Scheme 1. The closed state traps the nucleotide and aligns catalytic residues to facilitate fast catalysis. Although it is clear that Mg.dNTP is the substrate for the reaction, the roles of the second Mg^2+^ in each of these steps are not know and without this information the role of Mg^2+^ in specificity cannot be established. For example, what is the order of binding the second Mg^2+^ relative to other steps in the pathway? Does Mg^2+^ bind to the open state? What is the net *K*_*d*_ for binding the catalytic Mg^2+^ to either the open or the closed state? How does free Mg^2+^ ion affect ground-state binding, conformational change, chemistry, and PPi release? How does the free Mg^2+^ concentration alter nucleotide specificity? Is there evidence for the involvement of a third metal ion? In this study, we addressed these questions by examining the Mg^2+^ concentration dependence of each step in the reaction pathway in order to estimate the initial binding affinity of Mg.dNTP to the open state of the enzyme, the rate of the nucleotide-induced conformational change, the rate of nucleotide release before chemistry, the rate of the chemical reaction, and the rate of product release. This analysis allows us to resolve the contributions of each metal ion toward enzyme specificity. We also use MD simulations to view the binding of Mg^2+^ to multiple sites on the enzyme-DNA complex. Results from MD simulations are consistent with experimental measurements of binding affinity and provide molecular details for aspects of Mg^2+^ binding that cannot be observed directly.

**Scheme 1.**
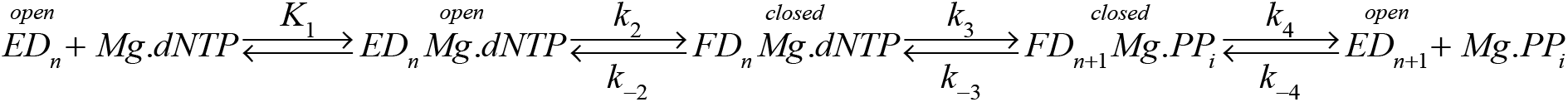
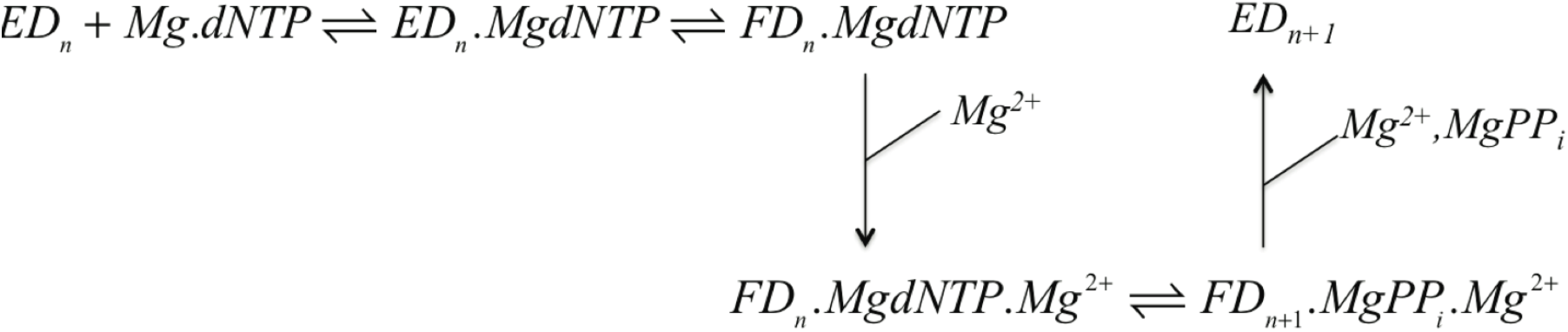
Pathway of DNA polymerization. The minimal reaction pathway is shown where *ED*_*n*_ and *FD*_*n*_ represent the enzyme-DNA complex in the *open* and *closed* states, respectively, as observed in crystal structures and shown to be kinetically important (15).

The studies performed here provide new insights toward understanding the role of metal ions in DNA polymerase fidelity. We show that Mg.dNTP is necessary and sufficient to induce the conformational change from the *open* to the *closed* state. The second Mg^2+^ binds after the conformational change, stabilizes the closed state, and stimulates the chemical reaction. Accordingly, we will refer the second metal ion as the *catalytic Mg^2+^* to distinguish it from the *nucleotide bound* Mg^2+^, although it must be clear that both metal ions are required for catalysis. In the course of performing these experiments, we also developed a simplified method to accurately define concentrations of free Mg^2+^ and Mg.dNTP in solution using a Mg-EDTA buffer.

## Results

To address the role of Mg^2+^ in catalysis and specificity, we systematically studied the effects of free Mg^2+^ concentration on each step of the nucleotide incorporation pathway outlined in Scheme 1: ground-state binding (*k_1_*), forward and reverse rates of the conformational change (*k_2_* and *k*_*-2*_), chemistry (*k_3_*), and PPi release (*k*_*4*_). We began by measuring the rate and equilibrium constants governing nucleotide binding and enzyme conformational dynamics.

### Effect of free Mg^2+^ concentration on Mg.dTTP binding kinetics and equilibrium

To study the kinetics and equilibria for binding of Mg.dTTP to HIVRT, we used HIVRT labeled with MDCC (7-diethylamino-3-((((2-maleimidyl)ethyl)amino)carbonyl) coumarin) on the fingers domain as described previously. The labeling provides a signal to measure the conformational changes between open to closed states (13). The fluorescence change was recorded using a stopped flow instrument after rapidly mixing various concentrations of Mg.dTTP with an enzyme-DNA complex formed with a primer that is dideoxy-terminated (ED_dd_) so that Mg.dTTP binds but does not react, as seen in crystal structures (17). The fluorescence signal is due to the fast closing of the enzyme after Mg.dTTP binding, but at low concentrations of nucleotide the rate is limited by the kinetics of binding, affording measurement of the net second-order rate constant for Mg.dNTP binding (Scheme 2).

**Scheme 2.**
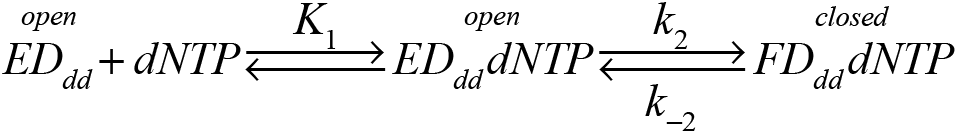
Two-step nucleotide binding. The simplified model shows only the binding and conformational change steps when chemistry is blocked by using a dideoxy-terminated DNA primer.

Under conditions of rapid equilibrium substrate binding, the reaction follows a single exponential with an observed decay rate (eigenvalue, λ) that is a hyperbolic function of the substrate concentration. At low substrate concentration, the slope of the concentration dependence defines the apparent second order rate constant for substrate binding, *k*_*on*_.

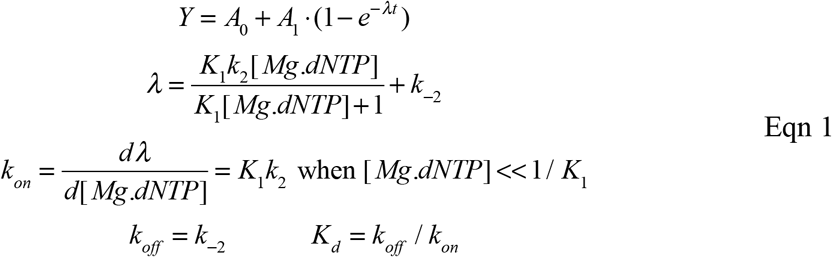

Because binding data were collected only at low nucleotide concentrations, the data did not resolve *k_1_* and *k_2_*; rather, we only determine the apparent second order rate constant for Mg.dNTP binding given by the product *k_1_k_2_*.

The kinetics of binding are shown at various concentrations of Mg.dTTP in Figures 1A and 1C in the presence of 10 mM and 0.25 mM free Mg^2+^, respectively. It should be noted that the *K*_*d*_ for Mg^2+^ binding to nucleotide is 29 μM (5), so the Mg.dTTP complex is saturated even at the lowest concentrations of Mg^2+^ used in this study. In Figures 1A and 1C, the rate and amplitude of the reaction increase as a function of Mg.dTTP concentration so that both the forward and reverse rate constants can be defined from the data. That is, the rate of binding is a function of the sum of the forward and reverse rate constants, while the fluorescence endpoint defines the equilibrium constant, which gives the ratio of the rate constants. In global data fitting, the information is combined to define both forward and reverse rate constants (*k*_*on*_ = *k_1_k_2_* and *k*_*off*_ = *k*_*-2*_). As a further test, we performed equilibrium titrations (Figures 1B and 1D for 10 and 0.25 mM Mg^2+^, respectively). Global fitting of the two experiments at each Mg^2+^ concentration using the model shown in Scheme 2 allows us to accurately define the net dissociation constant (*K*_*d*_ = *k*_*off*_/*k*_*on*_ = 1/(*k_1_k_2_*)) for Mg.dTTP as well as the rate constants governing binding and dissociation as summarized in Table 1. From these results we conclude that the free Mg^2+^ concentration has little effect on the net nucleotide binding affinity. In studies described below we resolve these two constants (*k_1_* and *k_2_*) using the full reaction with a normal DNA primer. The experiments reported here correlate our fluorescence signal with published structures and serve as a control for the effects of the dideoxy-terminated primer on the binding kinetics. In the next set of experiments, we measure the kinetics of reaction with a normal primer.

**Figure 1.**
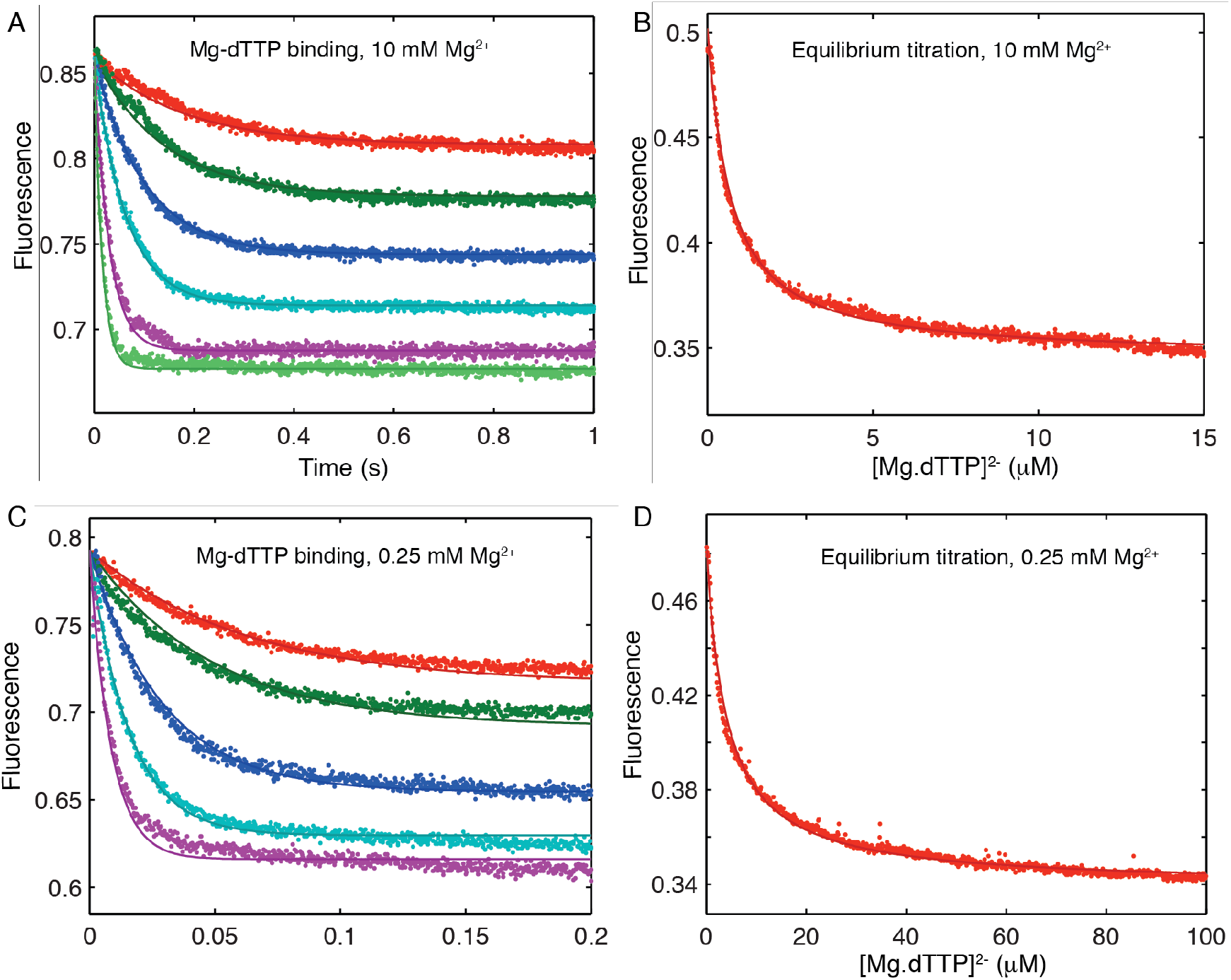
Kinetics of Mg.dTTP binding. The binding of Mg.dTTP was measured by stopped-flow fluorescence using a dideoxy-terminated primer to prevent chemistry (A). The experiments were performed by rapidly mixing various concentrations (0.5, 1, 2, 5, 10, and 20 μM) of Mg.dTTP with a preformed enzyme-ddDNA complex (100 nM MDCC-labeled HIV-1 wild type RT and 150 nM DNA with a dideoxy-terminated primer) in the presence of 10 mM free Mg^2+^. The binding of Mg.dTTP to HIVRT was also measured by equilibrium titration (B). The experiment was performed by titrating the preformed enzyme-DNA complex with varying concentrations of Mg.dTTP ranging from 0 to 20 μM. Global fitting of two experiments simultaneously to the model shown in Scheme 2 allows us accurately define the binding affinity and rate constants for association and dissociation of Mg.dTTP. The experiments in A and B were repeated in the presence of 0.25 mM Mg^2+^ to give the results shown in panels C and D. The results of data fitting are summarized in Table 1.

**Table 1.** Kinetic and equilibrium constants for Mg.dTTP binding to ED_dd_. Rate and equilibrium constants were derived in globally fitting data in Figure 1, according to Scheme 2.

| [Mg<sup>2+</sup>]<br>mM | $K_d$<br>(μM) | $k_{on}$<br>(μM<sup>-1</sup>s<sup>-1</sup>) | $k_{off}$<br>(s<sup>-1</sup>) |
| --- | --- | --- | --- |
| 10 | 0.67 ± 0.01 | 6 ± 0.1 | 4 ± 0.06 |
| 0.25 | 1.2 ± 0.07 | 8.8 ± 0.5 | 9.7 ± 0.2 |

### Complete kinetic analysis at 0.25, 1 and 10 mM free Mg^2+^

To measure the effects of free Mg^2+^ concentration on each step of the nucleotide incorporation pathway, experiments to measure the kinetics of nucleotide binding and dissociation, enzyme conformational changes, chemistry, and PPi release were all fitted globally at each free Mg^2+^ concentration. Figures 2, 3 and 4 show the results obtained at 10, 1 and 0.25 mM free Mg^2+^, respectively. Figure 2A shows the time dependence of the fluorescence change after mixing the enzyme-DNA complex with various concentrations of Mg.dTTP. In each single turnover experiment, the decrease in fluorescence is due to enzyme closing after nucleotide binding, while the increase in fluorescence is due to enzyme opening after chemistry.

**Figure 2.**
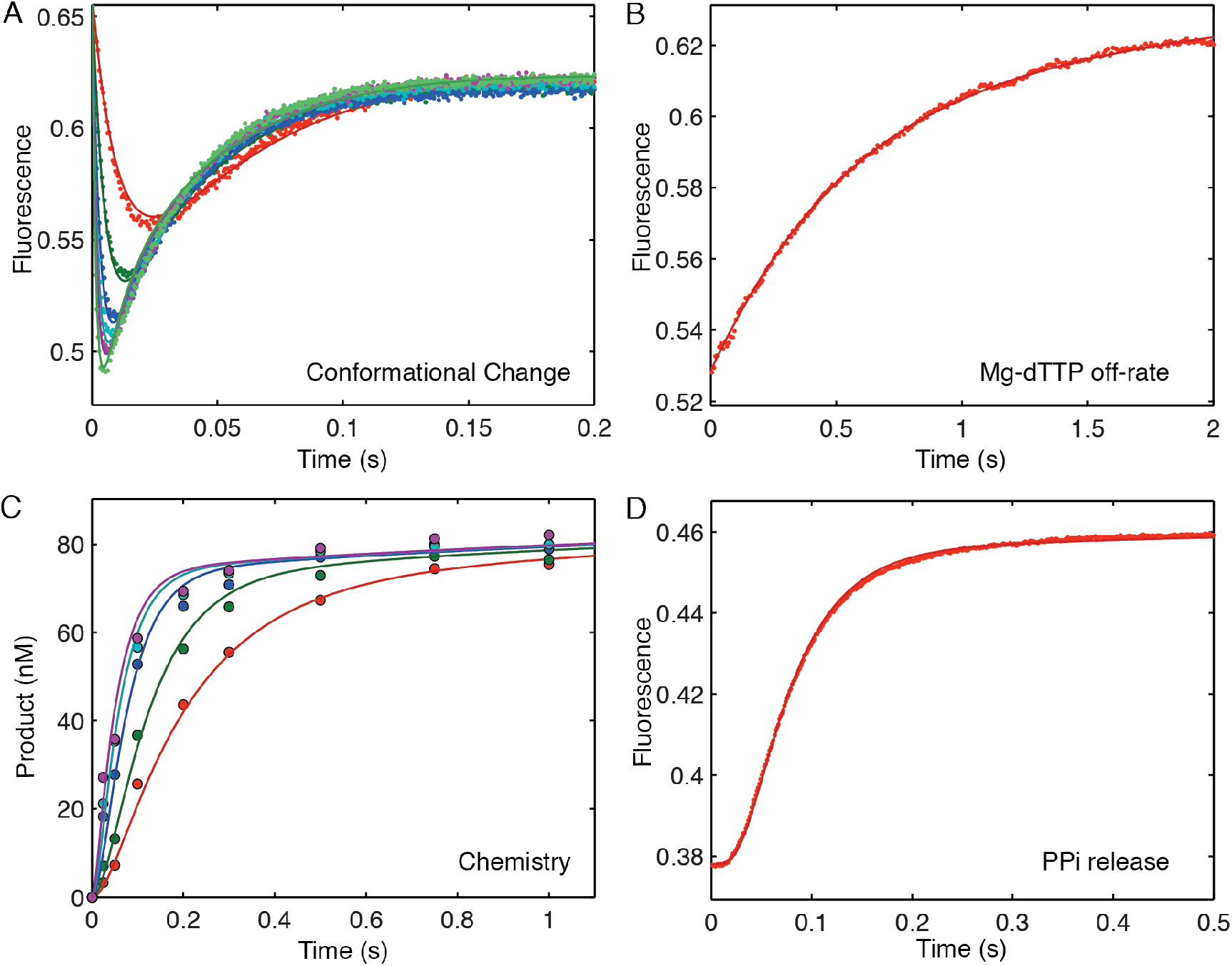
Correct nucleotide binding and incorporation in the presence of 10 mM Mg^2+^. (A) The time dependence of the fluorescence change upon dTTP incorporation was monitored by stopped flow fluorescence. The experiment was performed by rapidly mixing preformed ED complex (100 nM) with various concentrations (10, 25, 50, 75, 100, and 150 μM) of dTTP. (B) The nucleotide dissociation rate was measured by rapidly mixing preformed enzyme-DNAdd-dNTP complex (100 nM EDdd complex, 1 μM nucleotide) with a nucleotide trap consisting of 2 μM unlabeled ED complex and the fluorescence change was recorded to measure dNTP release. (C) The rapid chemical quench flow experiment was performed by rapidly mixing preformed ED complex (100 nM) with various concentrations (1, 2, 5, 10 and 20 μM) of dTTP. (D) The rate of PPi release was measured by a coupled pyrophosphatase/phosphate sensor assay (39). Four experiments were fit simultaneously to define the kinetic parameters governing nucleotide incorporation as shown in Scheme 1 and summarized in Table 2.

**Figure 3.**
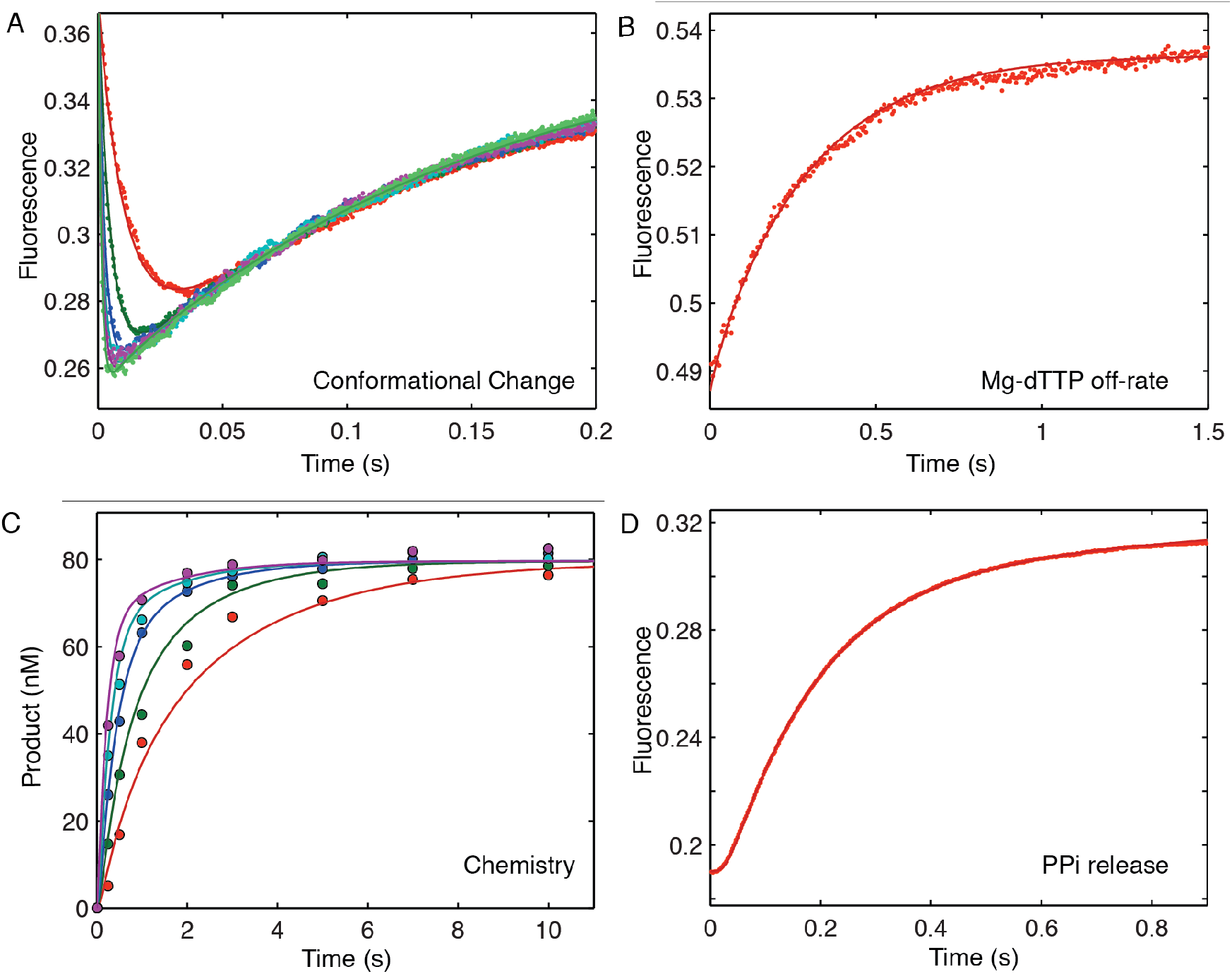
Correct nucleotide binding and incorporation in the presence of 1 mM Mg^2+^. Experiments were performed as described in Figure 2, but at 1 mM Mg^2+^. Four experiments were fit simultaneously to define each kinetic parameter governing nucleotide incorporation in the presence of 1 mM free Mg^2+^ as shown in Scheme 1 and summarized in Table 2.

**Figure 4.**
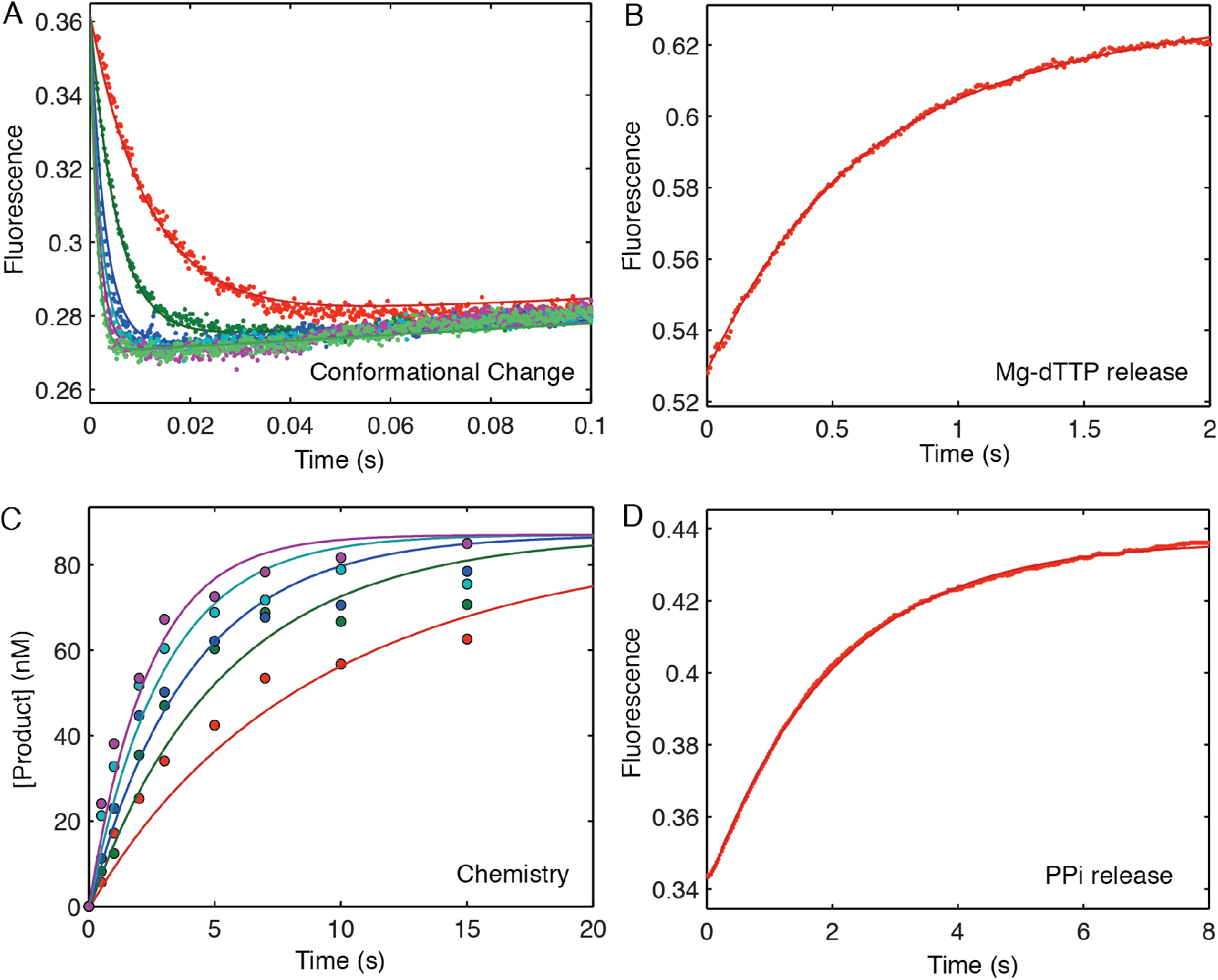
Correct nucleotide binding and incorporation in the presence of 0.25 mM Mg^2+^. Experiments were performed as described in Figure 2, but at 0.25 mM Mg^2+^, except that the method to measure nucleotide dissociation was modified due to the low efficiency of the nucleotide trap at low Mg^2+^ concentration. We used a combination of two enzymes (unlabeled HIVRT-DNA and apyrase) to trap and digest free nucleotides in solution (Figure 4B). The *k*_*-2*_ value obtained by this method was 9.9 ± 0.4 s^−1^. Four experiments were fit simultaneously to define each kinetic parameter governing nucleotide incorporation in the presence of 0.25 mM free Mg^2+^ as shown in Scheme 1 and summarized in Table 2.

**Table 2.** Kinetic constants for Mg.dTTP binding, conformational change, incorporation, and PPi release. Rate and equilibrium constants were derived in globally fitting data in Figures 2–5 as described in the Results. Rate constants refer to Scheme 1.

| [Mg<sup>2+</sup>]<br>(mM) | 1/$K_1$<br>(μM) | $k_2$<br>(s<sup>-1</sup>) | $k_{-2}$<br>(s<sup>-1</sup>) | $K_2$ | $k_3$<br>(s<sup>-1</sup>) | $k_4$<br>(s<sup>-1</sup>) |
| --- | --- | --- | --- | --- | --- | --- |
| 10 | 275 ± 3 | 2000 ± 67 | 3.9 ± 0.1 | 513 ± 22 | 21 ± 0.1 | > 100 |
| 1 | 213 ± 3 | 1960 ± 173 | 9.4 ± 0.2 | 209 ± 19 | 6 ± 0.1 | > 160 |
| 0.25 | 216 ± 2 | 1910 ± 178 | 9.7 ± 0.2 | 197 ± 19 | 0.6 ± 0.01 | > 20 |

To measure the rate of the reverse of the conformational change, the nucleotide dissociation rate was measured by rapidly mixing a preformed enzyme-DNAdd-dTTP complex (100 nM MDCC-labeled HIVRT, 150 nM 25ddA/45-nt, 1 μM Mg.dTTP) with a nucleotide trap that is consisting of 2 μM unlabeled enzyme-DNA complex (Figure 2B). Because the DNA primer was dideoxy-terminated, premixing the ED_dd_ complex with incoming nucleotide allows the binding but not the chemistry. After the addition of the large excess of the unlabeled enzyme-DNA complex, the rate of the fluorescence change defines the rate for the reverse of the conformational change, allowing rapid release of the nucleotide. Because these data were fit using computer simulation and we included the known kinetics of the nucleotide binding to the enzyme-DNA nucleotide trap, it was not necessary to repeat this experiment at multiple concentrations of the nucleotide trap. These results define Mg.dNTP binding and release and agree with the measurement of the net equilibrium constant shown in Figures 1B and D.

In the next experiment, the rate of chemistry was measured in a rapid quench-flow assay (Figure 2C). An enzyme-DNA complex (^32^P-labeled primer) was rapidly mixed with varying concentrations of incoming nucleotide. The rate of product formation as a function of nucleotide concentration was used to define the maximum polymerization rate (*k*_*cat*_) of the reaction and the specificity constant (*k*_*cat*_/*K*_*m*_).

To measure the rate of PPi release, a coupled pyrophosphatase/phosphate sensor assay was performed as described previously (18,19). Because there is a large fluorescent change upon phosphate binding to MDCC-PBP (MDCC–labeled phosphate binding protein) and the rate of phosphate binding to MDCC-PBP is much faster than that of the PPi release from HIVRT (20), the time course of the fluorescent change defines the rate of PPi release (18). Our data show that the rate of PPi release was coincident with the rates of chemistry and of reopening the enzyme as measured by the signal from fluorescently labeled HIVRT. In modeling the data by computer simulation, a lower limit on the rate of the PPi release was set at > 5-fold faster than the rate of the chemistry (Table 2), assuming opening was followed by PPi release. In fitting the data by simulation, the minimum rate of PPi release was defined as the value sufficient to make the two processes appear to coincide. Of course, we do not know the order of PPi release and enzyme opening because the two signals occur at the same rate.

The four experiments were fit simultaneously to rigorously define kinetic parameters governing nucleotide incorporation (Scheme 1) to get the results summarized in Table 2. To investigate the effects of free Mg^2+^ concentration on each step of the pathway, similar experiments and analyses were repeated at 1 and 0.25 mM free Mg^2+^ as shown in Figures 3 and 4, respectively. The kinetic parameters are summarized in Table 2. The results show that Mg^2+^ concentrations (from 0.25 mM to 10 mM) do not significantly affect the ground state Mg.dNTP binding (*k_1_*) or the rate of the conformational change (*k_2_*) but greatly affect the rate of the chemistry (*k_3_*) and less so the reverse of the conformational change (or enzyme reopening) (*k*_*-2*_). Although it appears that Mg^2+^ concentrations affect the rate of PPi release, this reflects the rates of chemistry and our simulation only gives a lower limit and therefore no direct conclusion regarding PPi release rate was possible. Our data support the conclusion that PPi release was not rate-limiting at either of the Mg^2+^ concentrations examined.

### Free Mg^2+^ concentrations do not affect the rate of the conformational change step

The experiments shown in Figures 2, 3 and 4 are not sufficient to resolve the rate of the conformational change step because it is too fast to measure directly at 37°C. Therefore, we measured the concentration dependence of the fluorescence transient at several temperatures and then extrapolated the observed rate constant (*k_2_*) to estimate the value at 37°C. The enzyme-DNA complex was rapidly mixed with various concentrations of dTTP as described in Figures 2A, 3A, and 4A but repeated at temperatures of 5, 10, 18, and 25°C. At each dTTP concentration the fluorescence transient was biphasic and was fit to a double exponential function:

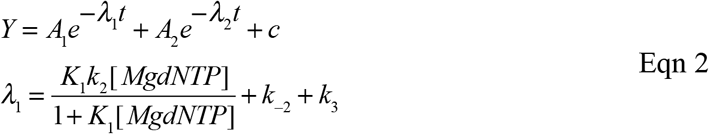

The concentration dependence of the rate of the fast phase of the fluorescence transient was then fit to a hyperbolic equation to obtain the maximum rate of the observed conformational change (λ_max_ = *k_2_ + *k*_*-2*_ + k_3_*) at each temperature. Note that *k*_*-2*_ and *k_3_* are much smaller than *k_2_* so λ_max_ ≅ *k_2_.* The experiments were repeated at various concentrations of free Mg^2+^: 0.25 mM (Figure 5A), 1 mM (Figure 5C), and 10 mM (Figure 5E). The observed maximum decay rate (λ_max_) observed at each temperature was then graphed on an Arrhenius plot (Figures 5B, 5D and 5F) and the values of *k_2_* at 37° at each of three different free Mg^2+^ concentrations were obtained by extrapolation by linear regression. The results indicate that Mg^2+^ concentrations (from 0.25 mM to 10 mM) do not affect the rate constant for the conformational change (*k_2_*) (Tables 2, 3). These results imply that Mg^2+^ binding from solution (at a concentration greater than 0.25 mM) is not required for the conformational change step, but it is required for the chemical reaction.

**Figure 5.**
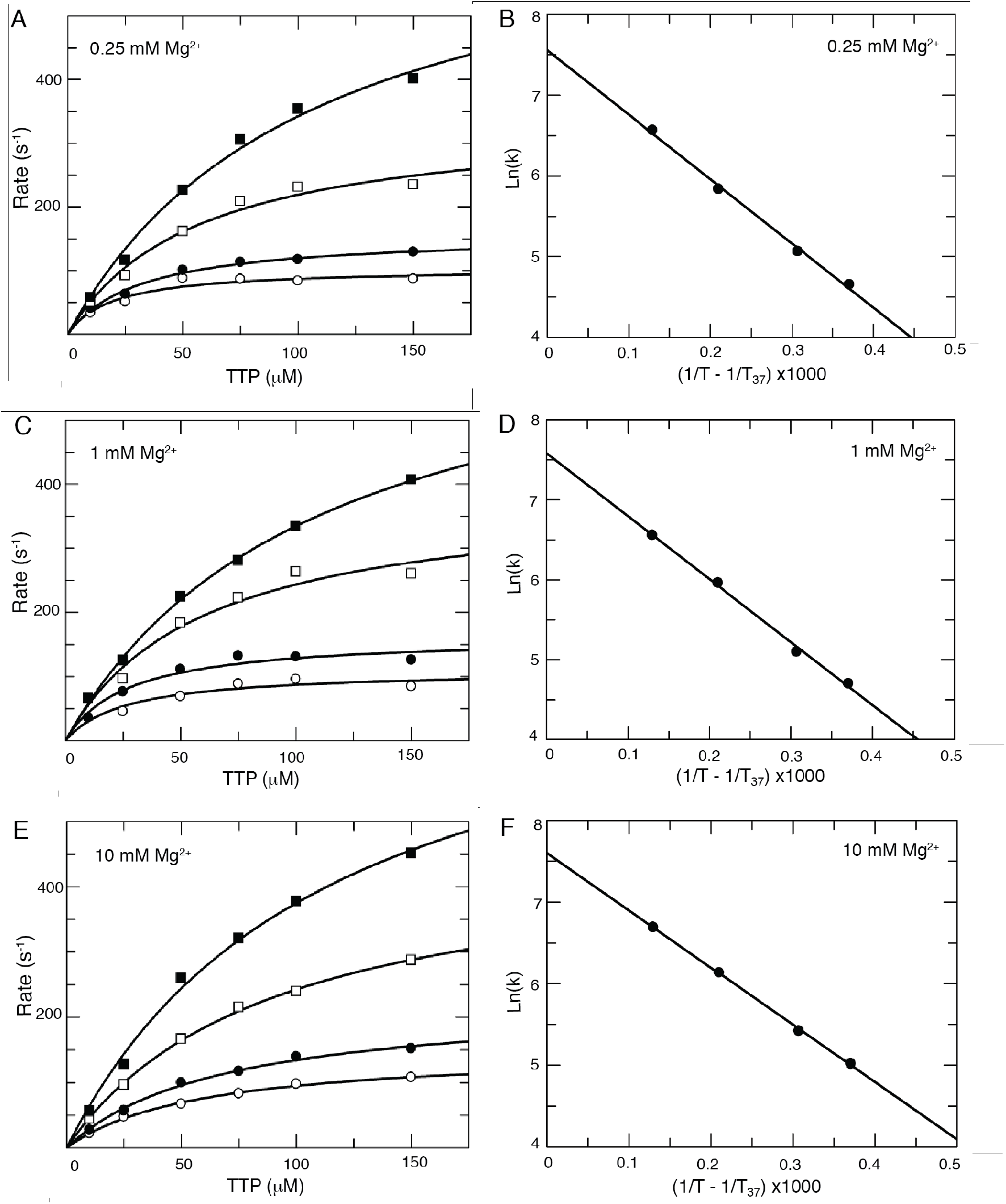
Mg^2+^ does not affect the rate of the conformational change. In order to estimate the rate of the conformational change at 37° C, the temperature dependence of the fluorescence change after dTTP binding was measured by stopped-flow methods at 0.25 mM (A and B), 1 mM (C and D) and 10 mM (E and F) free Mg^2+^ condition. At each Mg^2+^ concentration, an enzyme-DNA complex (50 nM) was rapidly mixed with various concentrations of dTTP (10, 25, 50, 75, 100 and 150 μM) at 5° (○), 10° (●), 18° (□), 25° (■) (panels A, C, and E). The concentration dependence of the rate of the fluorescence decrease upon nucleotide binding was fit to a hyperbolic equation to obtain the maximum rate of the conformational change (*k_2_*) at each temperature. The temperature dependence of *k_2_* was then analyzed on an Arrhenius plot (panels B, D and F) to estimate the maximum rate of the conformational change at 37° at each free Mg^2+^ concentration. The data show that the rate of the conformational change is independent of Mg^2+^ concentration (Tables 2, 3).

**Table 3.**
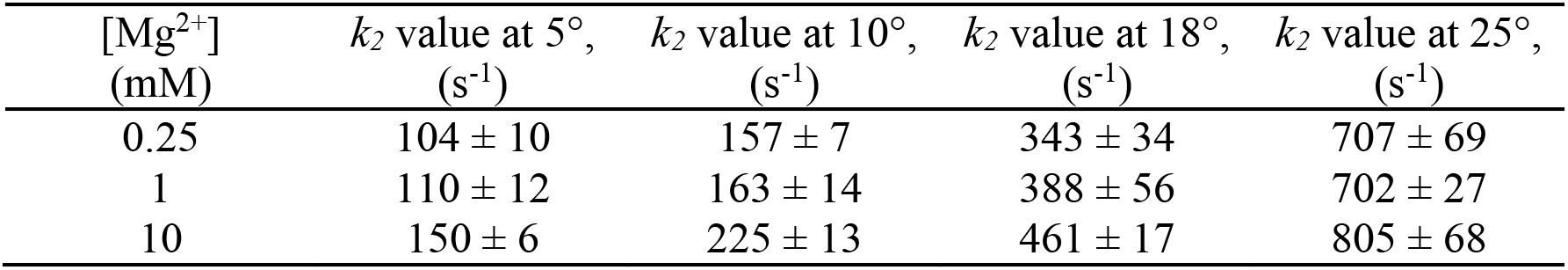
Temperature dependence of HIVRT conformational change at various concentrations of free Mg^2+^. Kinetic parameters were derived in fitting data in Figure 5 as described in the Results.

### The net K_d_ for Mg.dTTP binding

The substrate, Mg.dTTP binds initially to the open state with a relatively weak affinity, *K*_*d*_ = 275 μM at 10 mM Mg^2+^, which is followed by the conformational change leading to much tighter nucleotide binding. The net *K*_*d*_ for the two-step binding is defined by:

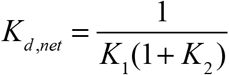

Accordingly, the net *K*_*d*_ for Mg.dTTP binding at 10 mM Mg^2+^ was 0.5 μM and increased to values of 1.0 and 1.1 μM at 1 and 0.25 mM Mg^2+^, respectively (Table 2). Thus, the nucleotide binding gets slightly tighter as the Mg^2+^ concentration increases, but this net effect is a product of opposing effects in the two-step binding, which we explore further below.

### Mg^2+^ concentration dependence of chemistry

The apparent binding affinity for the catalytic Mg^2+^ was measured more accurately by examining the Mg^2+^ concentration dependence of the rate of catalysis in a single turnover experiment. The experiment was performed by mixing an ED complex with solutions containing a fixed concentration of Mg.dTTP (150 μM) and various concentrations of free Mg^2+^ (ranging from 0.25 to 10 mM). This concentration of Mg.dTTP is much greater than its *K*_*m*_, so we are measuring the rate of the chemistry step in this experiment. The rates of chemistry, measured by rapid-quench and stopped-flow fluorescence methods, were observed for each reaction and plotted as a function of free Mg^2+^ concentrations (Figure 6A). The data were fit to a hyperbola to derive an apparent dissociation constant, *K*_*d,app*_ = 3.7 ± 0.1 mM, for the catalytic Mg^2+^. Because other known Mg^2+^ binding events reach saturation at much lower concentrations of Mg^2+^, they do not affect the measured *K*_*d*_ for the catalytic Mg^2+^. For example, the dissociation constant for the formation of the Mg.dTTP complex (28.7 μM) was 130-fold lower than the net *K*_*d*_ for binding of the catalytic Mg^2+^ (3.7 mM). Therefore, the observed concentration dependence of catalysis reflects only the binding of the catalytic Mg^2+^ to the closed ED-Mg.dTTP complex.

**Figure 6.**
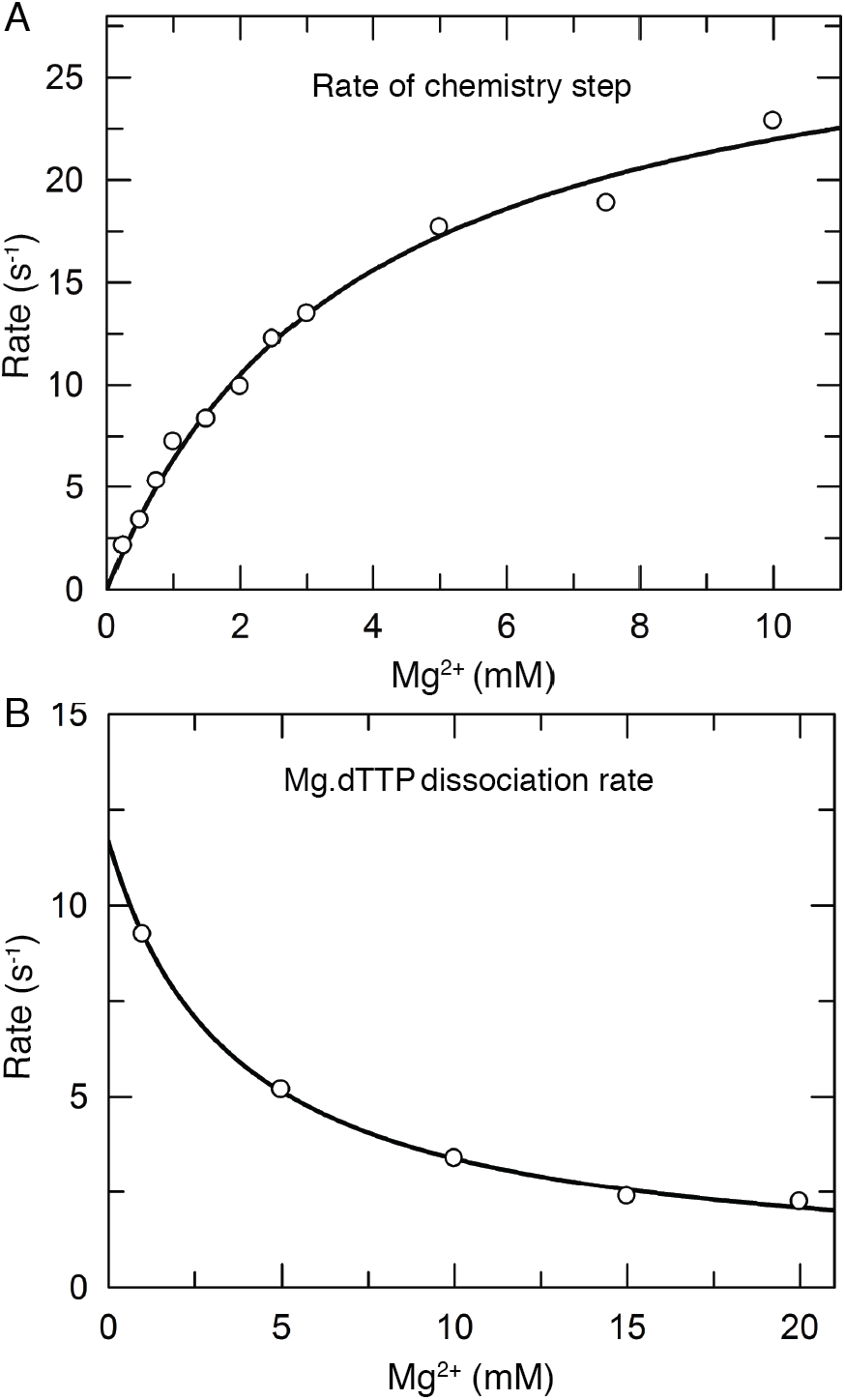
Mg^2+^ concentration dependence of the rates of chemistry and nucleotide release. The rate of polymerization was measured by mixing an ED complex with a fixed concentration of Mg.dTTP (150 μM) at various concentrations of free Mg^2+^, ranging from 0.25 to 10 mM. We then quenched the reaction with 0.5 M EDTA at various times and the amount of product formed was quantified so that the rate could be measured by fitting the time dependence to a single exponential. The measured rate was plotted as a function of free Mg^2+^ concentration, which was fit to a hyperbola to obtain the maximal rate of chemistry and the apparent dissociation constant (*K*_*d, app*_ = 3.7 ± 0.1 mM) for Mg^2+^ in stimulating the enzyme to catalyze the reaction.

### Binding of the Catalytic Mg^2+^ also affects the reverse of the conformational change

In Figure 6B we show the Mg^2+^ concentration dependence of the nucleotide dissociation rate, which we believe is limited by the rate of enzyme opening (*k*_*-2*_). We plotted the observed rate of nucleotide dissociation as a function of free Mg^2+^ concentration and fit the data to a hyperbola to provide an estimated *K*_*d*_ = 3.7 ± 0.2 mM. This defines the apparent *K*_*d*_ for the binding of Mg^2+^ to slow the rate of nucleotide dissociation. The Mg^2+^ dependence of the nucleotide dissociation rate parallels the concentration dependence of the observed rate of the chemical reaction (Figure 6A) suggesting that the two effects are due to the same Mg^2+^ binding event. The binding of the catalytic Mg^2+^ stabilizes the closed enzyme state as active site residues are aligned to carry out catalysis. While the effect of Mg^2+^ binding on the rate of chemistry is profound (from <0.1 to 25 s^−1^), there is only a modest decrease (~ 2-fold) in the rate of nucleotide dissociation as Mg^2+^ concentration is increased from 0.25 to 10 mM.

### Catalytic Mg^2+^ is not required for the enzyme closing

Our results imply that the Mg.dTTP alone is sufficient for nucleotide binding and enzyme closing because the rate of conformational change (*k_2_*) is independent of free Mg^2+^ concentration. To further test this postulate, a preformed ED_dd_ complex (100 nM MDCC-labeled HIVRT and 150 nM 25ddA/45nt DNA) was rapidly mixed with either 50 μM dTTP or Mg.dTTP. The change of fluorescence upon dTTP or Mg.dTTP binding was monitored by stopped flow methods. In each experiment, we added 50 μM dTTP, but the free Mg^2+^ concentration was controlled using EDTA to allow the formation of 50 μM Mg.dTTP or ~ 0 μM Mg^2+^ to give free dTTP. The results showed that Mg.dTTP but not dTTP induces the conformational change of HIVRT (Figures 7A and B). However, one could still argue that the trace of Mg^2+^ needed to form Mg.dTTP could influence the observed conformational change kinetics. As a further test, we examined the kinetics of the fluorescence change after adding Rh-dTTP, an exchange-inert metal-nucleotide complex. In the absence of free Mg^2+^, 50 μM Rh-dTTP induced a decrease of the fluorescence (Figure 7C) although slightly lower in rate and amplitude when compared to Mg.dTTP. These results indicate that the metal-nucleotide complex is sufficient to induce enzyme closing. However, the lower rate and amplitude seen with Rh-dTTP compared to Mg.dTTP reveals differences between the two metal ion complexes. These results support conclusions derived from analysis in Figures 2–5 (summarized in Tables 2 and 3) suggesting that the concentration of free Mg^2+^ does not alter the rate of the conformational change step.

**Figure 7.**
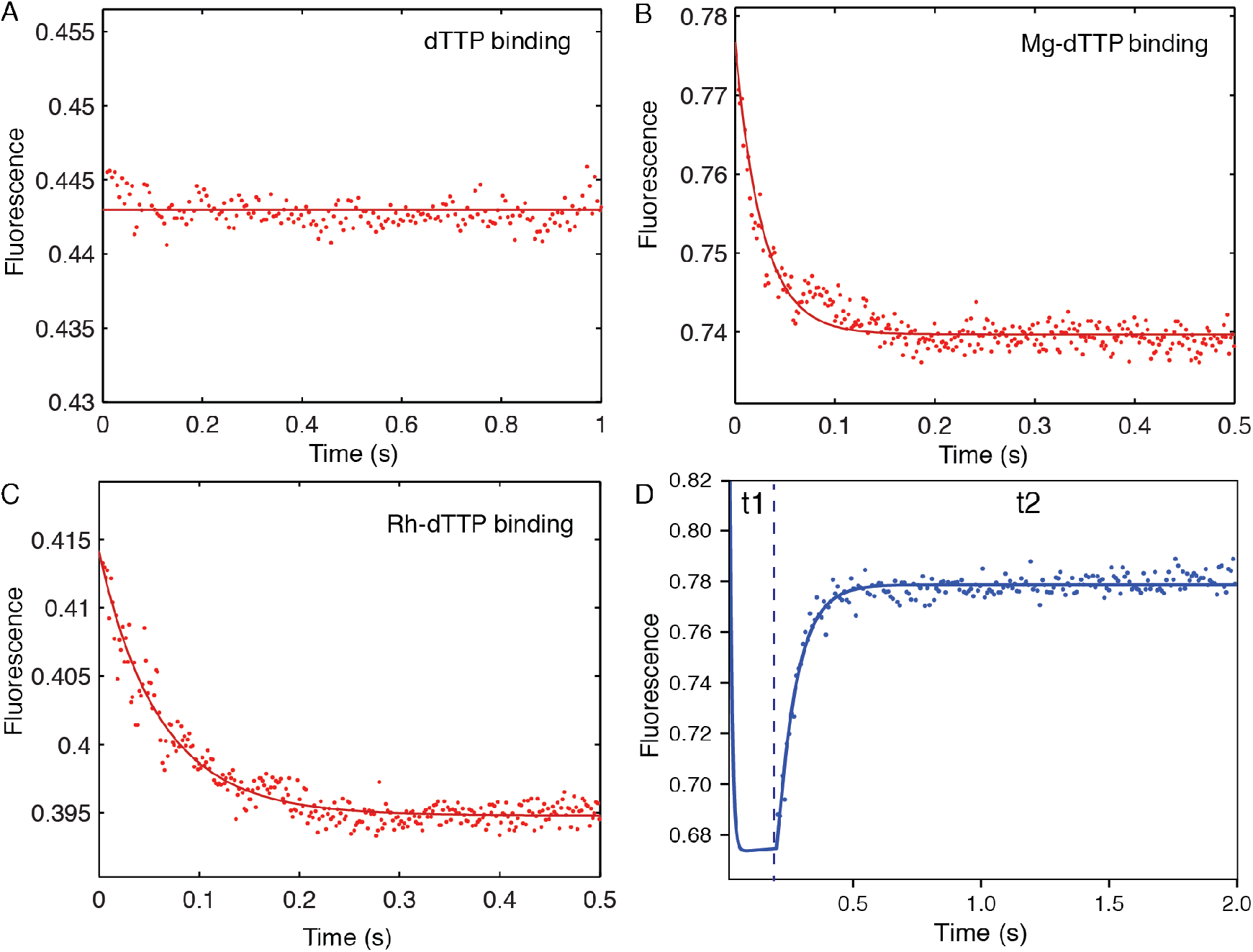
Role of Mg^2+^ in nucleotide binding-induced conformation change and catalysis. The role of Mg in the enzyme nucleotide-induced conformational change was examined using MDCC-HIVRT fluorescence under various conditions with a preformed EDdd complex (100 nM). (A). We rapidly mixed EDdd with dTTP (50 μM) in the absence of Mg^2+^ (concentration of EDTA: 500 μM). (B). We rapidly mixed EDdd with Mg.dTTP (50 μM) (concentrations of Mg.dTTP was controlled by using EDTA and simulated by using *KinTek Explorer* software). (C). We rapidly mixed EDdd with Rh-dTTP (50 μM) in the absence of Mg^2+^. (D). A double mixing experiment was performed by first mixing an ED complex (100 nM) with 10 μM Mg.dTTP in the presence of 25 μM free Mg^2+^ for 0.2s (t1), followed by second mixing with 10 mM Mg^2+^ (t2). The fluorescence increase after secondary mixing was monitored by stopped flow assay showing the fast opening of the enzyme after chemistry.

Because the catalytic Mg^2+^ binds relatively weakly, we can easily resolve effects of the metal ions on the conformational change versus chemistry by varying the Mg^2+^ concentration in the experiment. We also examined whether the catalytic Mg^2+^ participated in the conformational change by using a double mixing experiment (Figure 7D). The experiment was performed by first mixing the ED complex (100 nM MDCC-labeled HIVRT and 150 nM 25/45nt DNA) with 10 μM Mg.dTTP in the presence of 25 μM free Mg^2+^ for 0.2s (t1), followed by a second mixing with a large excess of free 10 mM Mg^2+^ (t2). The fluorescence change upon the second mixing was monitored by stopped flow fluorescence to measure the opening of the enzyme after chemistry. During the first mixing step at 25 μM free Mg^2+^ the half-life of dTTP incorporation was >4 s (Figure 2D); therefore, chemistry did not occur significantly during the first mixing step of 0.2 s. The apparent *K*_*d*_ = 3.7 mM predicts that only 0.7% of the catalytic Mg^2+^ binding sites will be occupied at 25 μM Mg^2+^. During the second mixing with the addition of a large excess of free Mg^2+^ (10 mM), the catalytic Mg^2+^ binds to HIVRT and stimulates catalysis, which is followed by rapid opening of the enzyme to give a fluorescence signal. These data demonstrate that Mg.dTTP is sufficient to induce enzyme closing without the catalytic Mg^2+^. If the catalytic Mg^2+^ were required for the closing of the enzyme, we would have observed a decrease in fluorescence followed by an increase after adding excess Mg^2+^ in the second mixing step. The immediate reopening of the enzyme directly after the second mixing (Figure 7D) demonstrates that Mg.dTTP alone is sufficient to induce the conformational change from the *open* to the *closed* state of HIVRT. The binding of the catalytic Mg^2+^ in the second mixing step is necessary for fast catalysis and enzyme opening but is not required for the conformational change step. The catalytic Mg^2+^ binds only after enzyme closing to stimulate catalysis.

### Mg^2+^ weakly competes with Mg.dTTP in binding to the open state of the enzyme

With a pair of aspartic acid residues in the active site, one might expect that Mg^2+^ could bind tightly to the open state of the enzyme in the absence of nucleotide, but this is never seen in crystal structures. We reasoned that if Mg^2+^ binds to the open state of the enzyme-DNA complex, then it could be a competitive inhibitor of Mg.dNTP binding. A slight effect can be seen in the data in Table 2. Values of the apparent *K*_*d*_ for Mg.dNTP in the ground state binding to the open form of the enzyme (1/*k_1_*) increase as the concentration of Mg increases. Although we have only three data points because of the extensive analysis required to derive this number, the data can still provide an estimate of the *K*_*d*_ for Mg^2+^ binding based on the observed competition according to the following relationship:

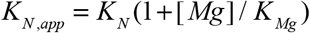

where *K*_*N*_ and *K*_*Mg*_ are the *K*_*d*_ values for Mg.TTP and Mg^2+^, respectively and *K*_*N,app*_ is the apparent *K*_*d*_ for nucleotide binding (given in Table 2). Linear regression of a plot of *K*_*N,app*_ versus [Mg] gives an estimate of *K*_*N*_ = 212 ± 4 μM and *K*_*Mg*_ = 34 ± 4 mM. This analysis suggests that the binding of Mg^2+^ to the open state of the enzyme in the absence of Mg.dNTP is tenfold weaker than binding to the closed state in the presence of Mg.dNTP.

### Mg^2+^ concentration effects on nucleotide specificity

The specificity constant (*k*_*cat*_/*K*_*m*_) defines fidelity for nucleotide incorporation in comparing a cognate base pair with a mismatch. The specificity constant is best understood as the second-order rate constant for substrate binding times the probability that, once bound, the substrate goes forward to form and release product. Steady state kinetic parameters were calculated from the primary rate constants (Eqn 2) to get the results summarized in Table 4. Our results showed that the value of the specificity constant is decreased by 12-fold as the free Mg^2+^ concentration is changed from 10 to 0.25 mM due to the slower rate of incorporation and change in specificity-determining step. In addition, the results in our study suggest that nucleotide specificity is re-defined as the free Mg^2+^ concentration is altered. Because the rate of the conformational change (*k_2_*) is much faster than chemistry (*k_3_*), the specificity constant depends on the kinetic partitioning governed by the relative values of *k*_*-2*_ versus *k_3_* (13,16). If *k*_*-2*_ ≫ *k_3_*, the ground state binding and conformational change come to equilibrium and the specificity constant is governed by the product of binding equilibria and the rate of chemistry (*k*_*cat*_/*K*_*m*_ = *k_1_k_2_k_3_*). If *k*_*-2*_ ≪ *k_3_*, the nucleotide binding fails to reach equilibrium during turnover and the rate of chemistry does not contribute to the specificity constant; rather it is defined only the binding and conformational change steps (*k*_*cat*_/*K*_*m*_ = *k_1_k_2_*). As Mg^2+^ concentration decreases, the rate of dissociation increases, while the rate of chemistry decreases. With the correct nucleotide (dTTP) incorporation at 0.25 mM Mg^2+^, the value of *k*_*-2*_ (9.7 ± 0.2 s^−1^) is much greater than that of *k_3_* (0.6 ± 0.01 s^−1^), suggesting that the *k*_*cat*_/*K*_*m*_ value is governed by the product of binding equilibrium constants and the rate of chemistry (*k*_*cat*_/*K*_*m*_ = *k_1_k_2_k_3_*). When the free Mg^2+^ concentration was increased to 10 mM, the value of *k*_*-2*_ (3.9 ± 0.1 s^−1^) is smaller than that of *k_3_* (21 ± 0.1 s^−1^) and therefore *k*_*cat*_/*K*_*m*_ is largely governed only by the nucleotide binding (*k*_*cat*_/*K*_*m*_ = *k_1_k_2_*). The mechanistic basis for nucleotide specificity changes as a function of the free Mg^2+^ concentration. We have not examined the kinetics of misincorporation at the lower Mg^2+^ concentrations.

**Table 4.** Steady state kinetic parameters versus concentration of free magnesium ion. The steady state and equilibrium constants were calculated as described in the Methods.

| [Mg<sup>2+</sup>]<br>(mM) | $K_{d,net}$<br>(μM) | $K_m$<br>(μM) | $k_{cat}$<br>(s<sup>-1</sup>) | $k_{cat}/K_m$<br>(μM<sup>-1</sup>s<sup>-1</sup>) | Fold change<br>in $k_{cat}/K_m$ |
| --- | --- | --- | --- | --- | --- |
| 10 | 0.5 ± 0.02 | 3.6 ± 0.1 | 20.7 ± 1 | 6 ± 0.3 | 1 |
| 1 | 1.0 ± 0.09 | 1.7 ± 0.2 | 6 ± 0.7 | 3.5 ± 0.6 | 0.6 |
| 0.25 | 1.1 ± 0.1 | 1.2 ± 0.1 | 0.6 ± 0.08 | 0.5 ± 0.08 | 0.08 |

We can illustrate the effects of free Mg^2+^ on free energy profiles of governing nucleotide incorporation at three different free Mg^2+^ concentrations (0.25 mM, 1 mM, and 10 mM) as shown in Figure 8. The *k*_*cat*_/*K*_*m*_ value is determined by the energy barrier between its highest peak relative to its unbound state. At 0.25 mM free Mg^2+^, the highest peak is the state between FD_n_N and FD_n+1_PPi (or chemistry step). Therefore, nucleotide specificity is determined by all of the steps from its unbound state to the chemistry (*k*_*cat*_/*K*_*m*_ = *k_1_k_2_k_3_*). At 10 mM free Mg^2+^, the highest peak is the state between ED_n_N and FD_n_N (or conformational change step). Thus, nucleotide specificity is determined by only two steps including ground state binding and the conformational change (*k*_*cat*_/*K*_*m*_ = *k_1_k_2_*). At 1 mM free Mg^2+^, the highest peak is not obvious by inspection and therefore a simplified equation for defining nucleotide specificity cannot be applied. To accurately define the nucleotide specificity constant (*k*_*cat*_/*K*_*m*_), the complete equation (Eqn 2) containing each parameter has to be used. The free energy profiles showed that nucleotide specificity is re-defined as the free Mg^2+^ concentration is altered from 10 mM to 0.25 mM.

**Figure 8.**
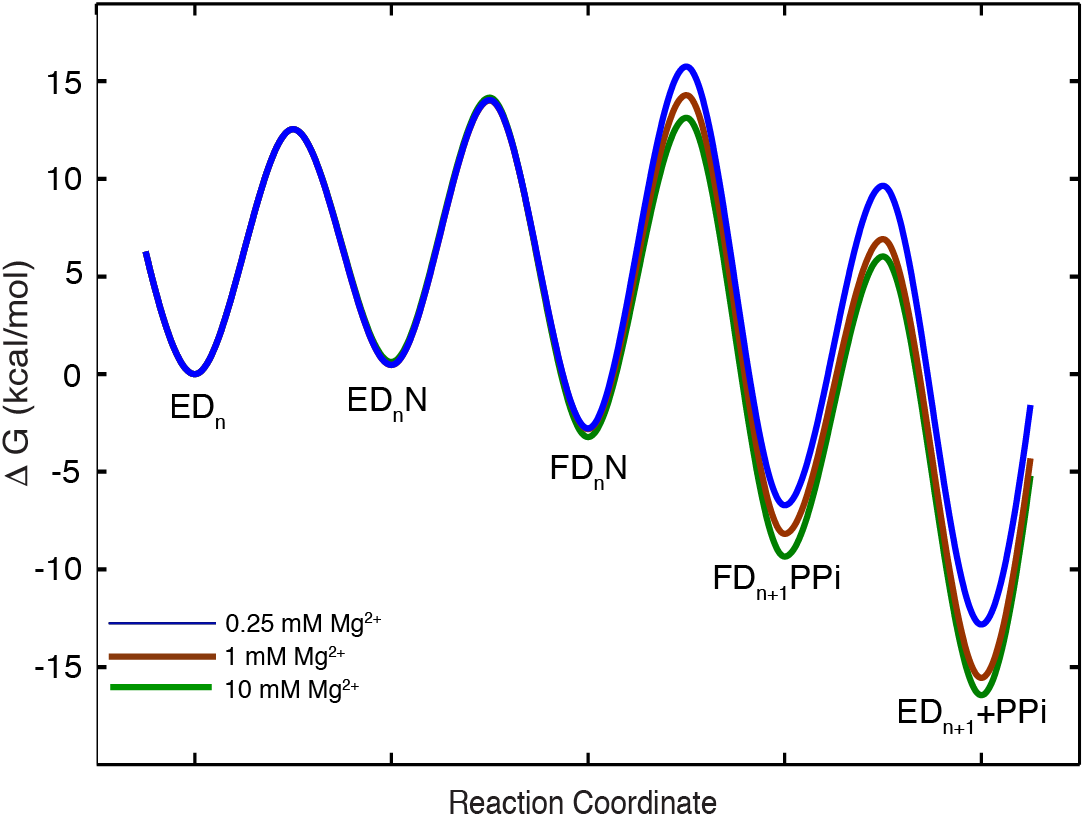
Free energy profile for nucleotide binding and catalysis at various Mg^2+^ concentrations. The free-energy diagrams for dTTP incorporation at 0.25, 1, and 10 mM free Mg^2+^ concentrations are shown in blue, brown and green colors. The free energy was calculated as ΔG = RT[ln(*k*T/h)-ln(*k*_*obs*_)] kcal/mol using rate constants derived from global fitting, where the constant k is the Boltzmann constant, T is 310 K, h is Planck’s constant, and *kobs* is the first-order rate constant for each step. The nucleotide concentration was set equal to 100 μM to calculate *kobs* as the pseudo-first order rate constant for nucleotide binding. Nucleotide binding to the open state (ED) was assumed to be diffusion-limited with *k_1_* = 100 μM^−1^s^−1^.

Because the *K*_*d*_ of catalytic Mg^2+^ may be lower than the intracellular concentration, our results suggest that binding of the catalytic Mg^2+^ provides the final checkpoint for nucleotide specificity as one component of the kinetic partitioning of the closed ED-Mg.dNTP complex so the substrate either dissociates or reacts to form products.

### Relating kinetics to available structures

In light of our kinetic results, we analyze the published structures with the focus on the coordination and interaction around Mg^2+^ ions. Magnesium prefers an octahedral coordination and will be most tightly bound when this geometry is satisfied (Figure 9A). Mg^2+^ ions at the polymerase domain of HIV-RT are coordinated through polar interactions with the side chains of aspartates 110 and 185 and the three phosphates of the nucleotide substrate (Figures 9B & 9C). In the nucleotide bound Mg^2+^ (MgB) in Figure 9B-C), the carboxylate side chains of Asp110 and 185 as well as the two non-bridging oxygens from the phosphates of the nucleotide appear to form the four coordination on the plane with a distance around 2.2-2.4 Å. Another phosphate oxygen and the carbonyl oxygen of valine 111 are at the apex from the opposite sides with a distance close to 2.6 Å. Therefore, MgB displays classic octahedral coordination geometry (Figure 9B). On the other hand, the catalytic Mg^2+^ (MgA) in Figure 9B-C), deviates from the standard octahedral coordination with what appears as four coordination by the side chains of Asp110 and Asp185 but not forming a plane. The two apex coordination sites are occupied by nucleotide on one end but empty on the other (Figure 9C), which is presumably occupied by a solvent water molecule that is not seen in the structure. The analysis of the metal coordination indicates that the two magnesium ions are bound differently, at least as observed by the refined crystal structures.

**Figure 9.**
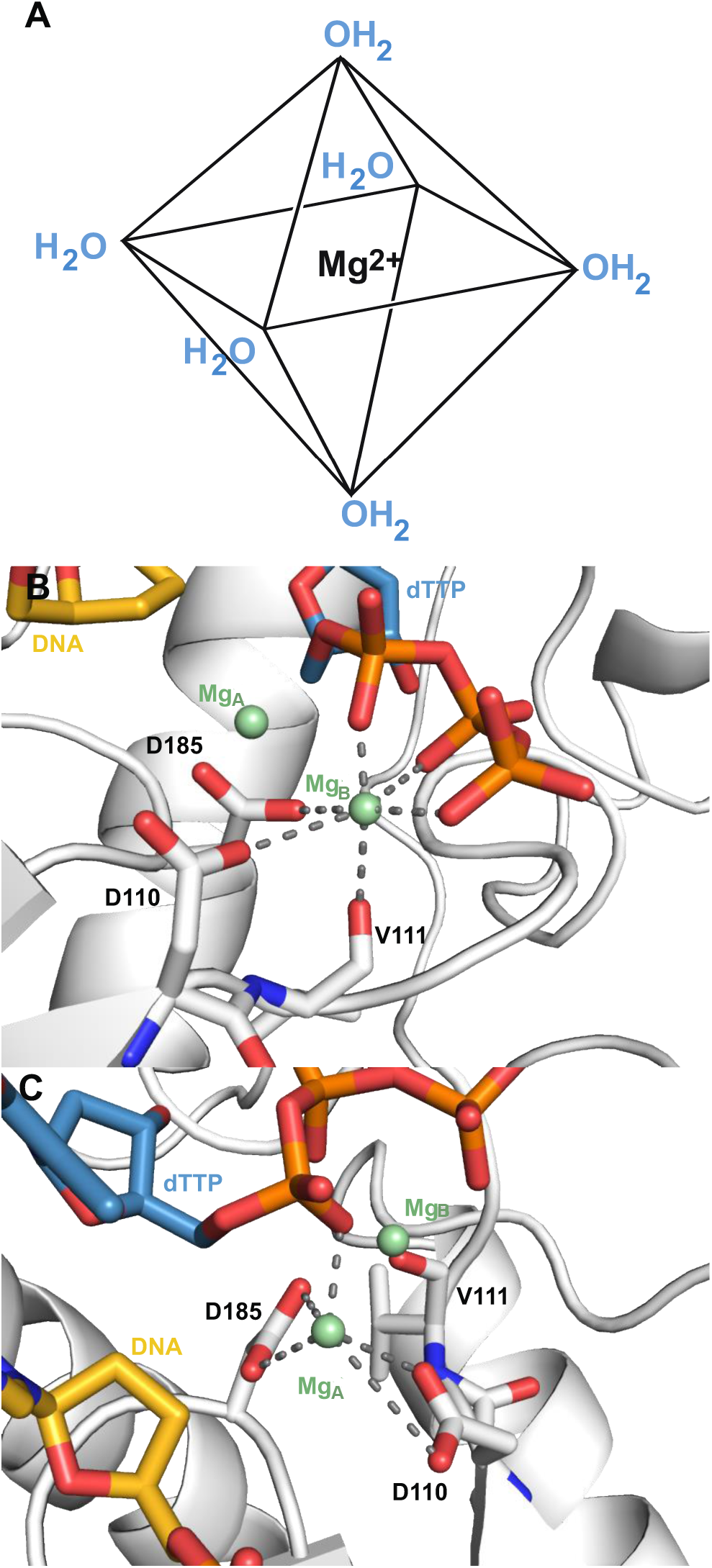
HIV-RT magnesium coordination geometry. A) Octahedral coordination geometry is ideal for magnesium ions. Magnesium preferentially forms six evenly spaced polar interactions, demonstrated here as interactions with solvent waters. B) Mg_B_ (pale green) at the HIV-RT polymerase domain exhibits ideal coordination geometry by forming three polar contacts with active site residues aspartate 110, 185 and valine 111(white), and three polar contacts with the triphosphate group of the substrate dTTP (steel blue) (PDB ID 1RTD). C) Mg_A_ (pale green) at the HIV-RT polymerase domain exhibits atypical coordination geometry and forms four polar contacts with the side chains of aspartate 110 and 185 (white) and one polar contact to the triphosphate group of substrates dTTP (steel blue) (PDB ID 1RTD).

To understand the relative occupancy and mobility of the ions, we scrutinized the temperature factors for each ion. Temperature factor (or thermal factor, B-factor) is defined as a measure of deviation of an atom from a certain position. A high B-factor correlates to high movement and low occupancy. Within the same molecule, the B-factor can be used to estimate the relative mobility of the atoms. The B-factors when both Mg^2+^ ions are present indicate that the mobility of the ions varies in HIV-RT. For the PDB ID 4PQU in which crystals were formed at high Mg^2+^ concentration (10 mM), MgA and MgB exhibit comparable B-factors (30.69 vs. 35.75, Å^2^ respectively). In other crystallization conditions in which HIV-RT was examined at lower concentrations of Mg^2+^ or with low-resolution diffraction, only one Mg^2+^ can be modeled in the density, which is consistently MgB (Table 5). In particular, MgA is poorly coordinated and displays a high relative B-factor or is missing in the structure. These structural data are in line with the measurement of the weak binding of *K*_*d*_ ≈3.7 mM for the catalytic Mg^2+^ observed in our solution study. In addition, no strong Mg^2+^ density was observed in the structure PDB ID 3KJV in spite of the high Mg^2+^ ion concentration of 10 mM. In this structure, HIV-RT is complexed with DNA:DNA primer/template only, consistent with our estimate of very weak binding of Mg^2+^ to the open ED complex without the Mg.dNTP. This analysis provides a structural evidence to support our kinetic analysis concluding that Mg.dNTP binds first to induce enzyme closing and the catalytic Mg^2+^ binds weakly and is only seen in the closed state.

**Table 5.**
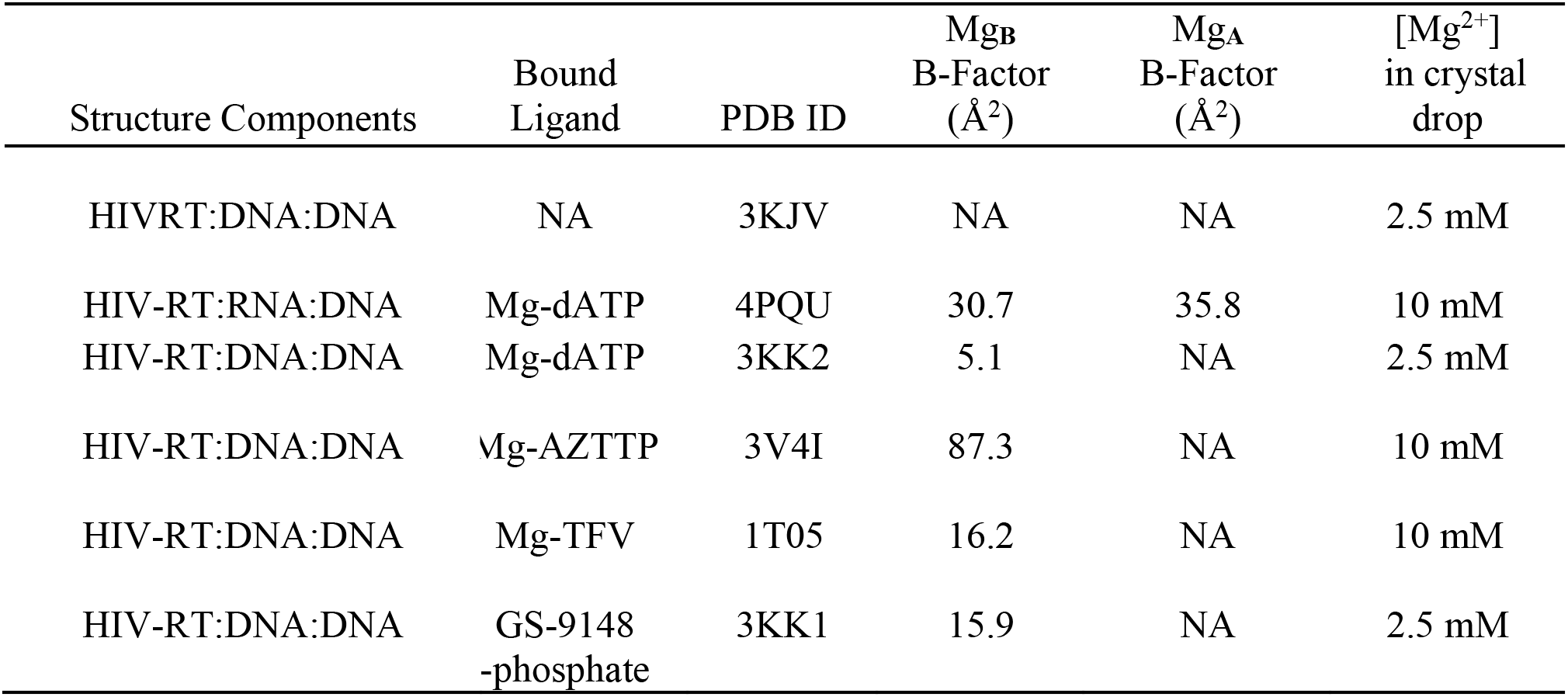
Analysis of HIVRT Crystal Structures. Published structures (cited in the table) were analyzed to quantify the binding of the nucleotide-bound Mg and catalytic Mg (MgA and MgB, respectively, as labeled in Figure 9)

**Table 6.** Kinetics of catalytic Mg ion binding and dissociation from all atom computer simulation study. The process is divided into binding from solution to carboxylic acid oxygens of 110 and 185 followed by exchange of two water from solvation shell of Mg^2+^(H_2_O) with the carboxylic acid oxygens.

| $k_1$<br>($M^{-1}s^{-1}$) | $k_{-1}$<br>($s^{-1}$) | $k_2$<br>($s^{-1}$) | $k_{-2}$<br>($s^{-1}$) | $1/K_1$<br>(mM) | $K_2$ |
| --- | --- | --- | --- | --- | --- |
| $7.6 \times 10^7$ | $3.2 \times 10^6$ | $1.6 \times 10^7$ | $2.3 \times 10^4$ | 42 | 700 |

### MD simulations to refine our understanding

Despite the significant evidence from kinetic analysis and structural data listed above suggesting a weaker binding and more rapid exchange of the catalytic Mg (MgA), our solution study is lacking atomic details. Structural studies provide valuable information at the atomic level but do not provide kinetic and thermodynamic parameters that define the role of Mg^2+^. In addition, it is naïve to think that Mg^2+^ only binds to the tight-binding, static sites seen in the crystal structures as there are many electronegative sites available to attract positively charged ions. To fill this void and complement our kinetic studies, we performed MD simulations to gain further insights into metal ion coordination along the pathway of DNA polymerase. Both matched and mismatched nucleotides are studied to understand the role of metal ions in the enzyme’s function. Computer simulations of Mg^2+^ coordination provide molecular level details that we cannot measure directly. We checked the validity of the MD simulations by comparison with what we can measure; namely, Mg^2+^ binding affinities in the open and closed states, before and after Mg.dNTP binding, respectively.

Figure 10 shows the distribution of Mg^2+^ ions around the enzyme-DNA-Mg.dNTP ternary complex, where each dot represents a Mg^2+^ position from simulation. To show the correlation of the positions sampled by Mg^2+^ ions we combine snapshots taken every 1 ns to give a visual image, reflecting the probability density distribution. The most obvious conclusion of this analysis is that there is a dense cloud of Mg^2+^ counterions surrounding exposed DNA and exposed charges on the surface of the protein. To further quantify the cation distribution, we plot the constant density regions as a heat map (Figure 11A). Details of how the ion densities are computed can be found in the Methods section. Here, the yellow regions show local Mg^2+^ concentrations in the range of 2-10 M, while the regions colored in red represent concentrations greater than or equal to 40 M. Simulations suggest a cloud of Mg^2+^ counter-ions surrounds the exposed DNA, as described by Manning theory (21), giving rise to an average local concentration of [Mg^2+^] ≈ 2.8 M near the DNA with an average number of bound Mg^2+^ ions of *N*_*Mg*^2+^_ ≈ 9.8. Note that we performed the MD simulation using a bulk solution concentration of 30 mM to have a statistically significant number of ions in simulation box. However, the excess cations surrounding the duplex is expected to be relatively insensitive to the bulk solution concentration (22).

**Figure 10.**
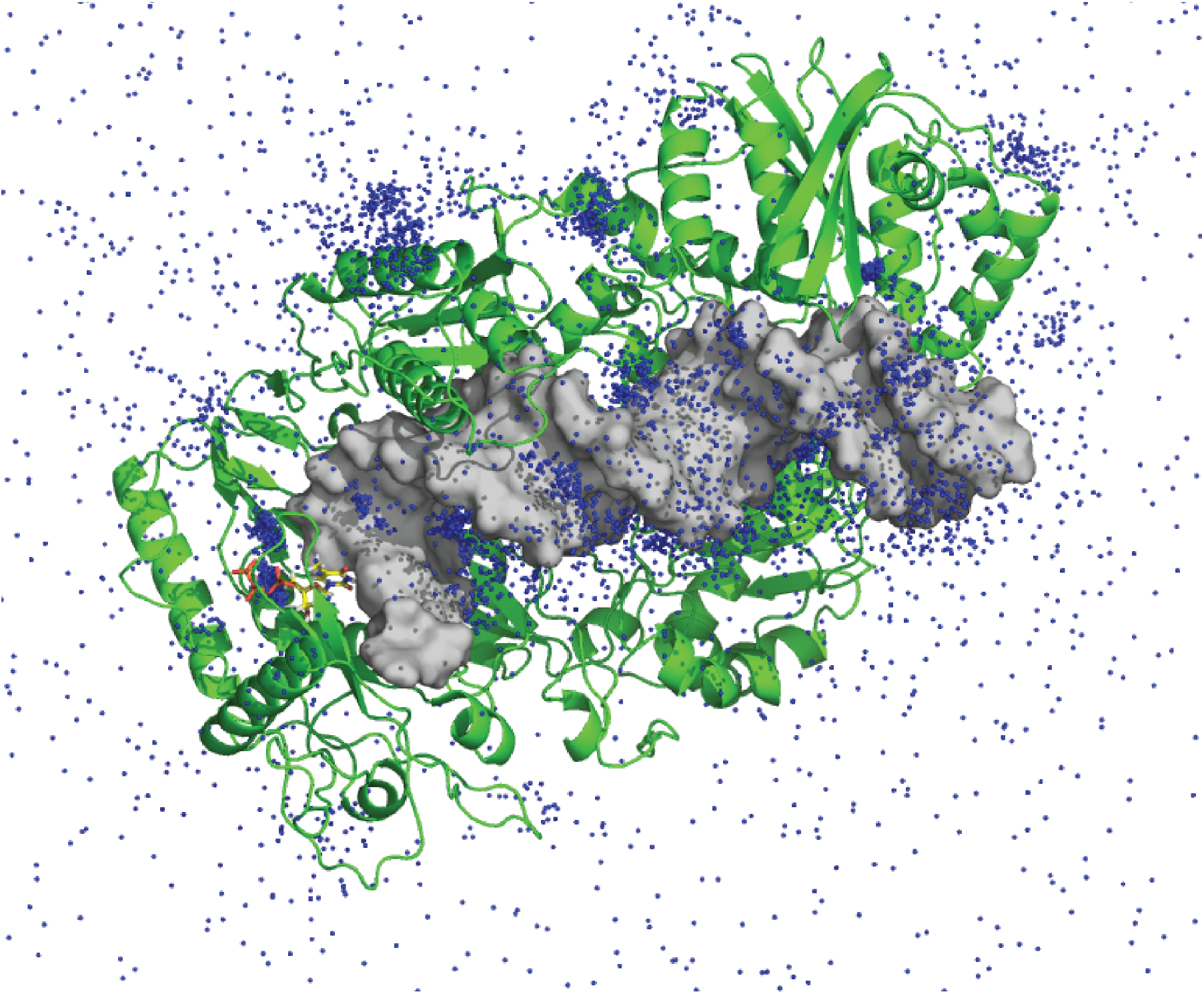
Mg ion distribution from simulations. Mg^2+^ ion positions sampled every 1 ns are combined and represented as dots to give a visual image of the probability density distributions sampled during a 300 ns molecular dynamics simulation.

**Figure 11.**
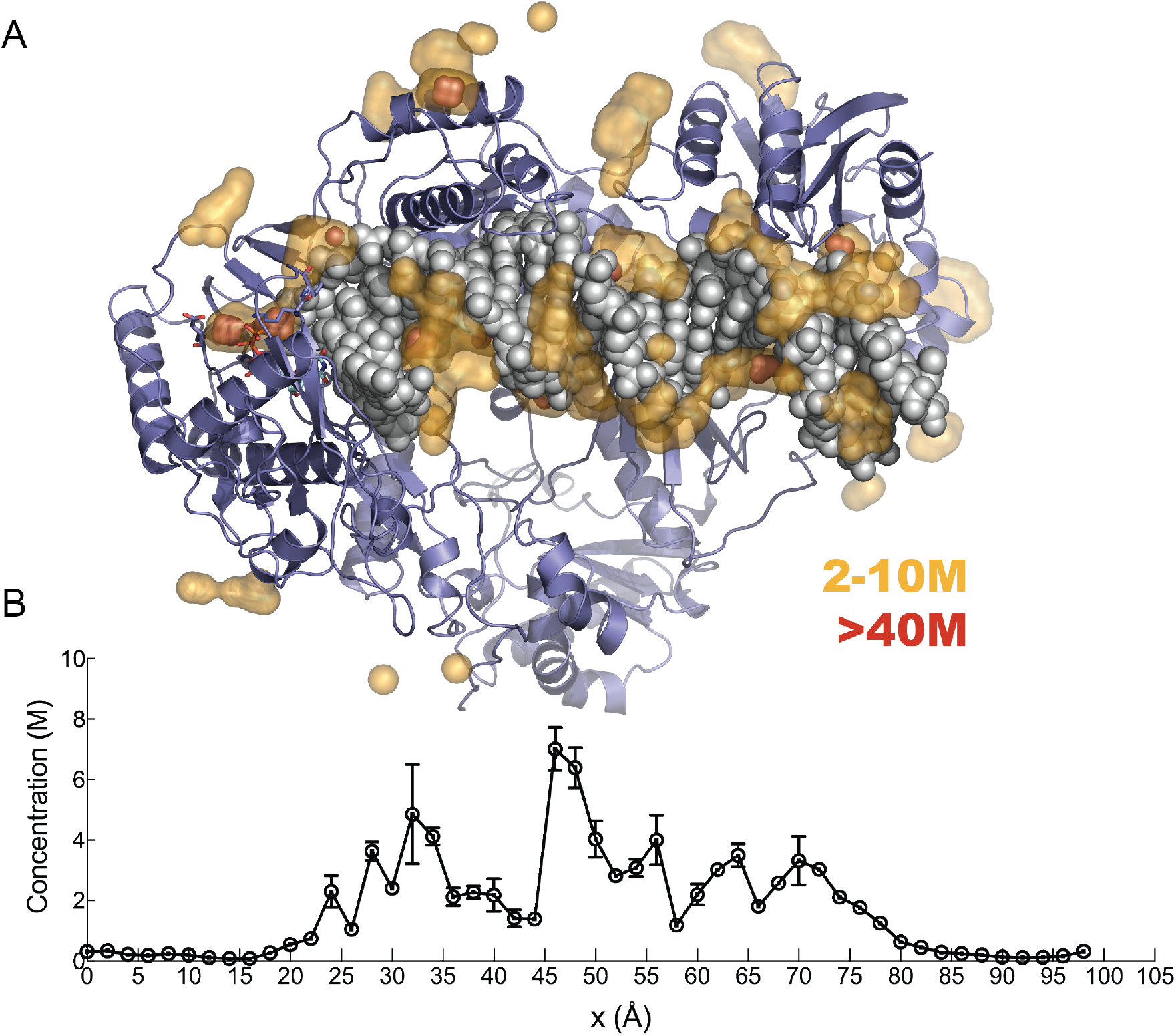
**A.** Overview of the Mg ion occupancy around the HIVRT. Local concentrations shown in a heatmap sticks represent the incoming nucleotide and two aspartic acid side chains. Protein is shown in cartoon representation while DNA in sphere. **B.** Mg ion density profile along the long DNA axis. The two images aligned for a clear view of the positions of the localized ions.

In Figure 11B, we project the local concentration of Mg^2+^ ions along the axis of the DNA helix. To compute the local cation concentration around DNA, we create a cylindrical shell of 15 Å radius covering DNA. Interestingly, the average concentration of Mg^2+^ ions around the DNA is not uniform along the DNA axis (Fig 11B); it is highest at the center of the DNA where DNA is most exposed to the solvent. Regions of lower concentration at 35-45Å and 55-65 Å coincide well with the thumb-site and RNase H domains, respectively. The positively charged residues at these domains create a depletion zone for free Mg^2+^ ions. The important conclusion from these observations is that a significant density of Mg^2+^ counter-ions surrounds the DNA. These cations likely impact the binding of DNA as well as the translocation of DNA during processive polymerization. The counterion atmosphere is not seen in the crystal structures because they bind diffusively, but they are important, nonetheless.

In addition to the diffusively bound counterions, MD simulations identify the specifically bound Mg^2+^ ions in agreement with crystal structures as discussed in the previous section. The comparison of metal ion binding sites with crystal structures allows us to benchmark simulation results and to further extrapolate the metal ion coordination to functional states where crystal structures are not readily available. Figure 12 summarizes our results for Mg^2+^ ion coordination along the polymerase reaction pathway for matching and mismatching dNTP bound states.

**Figure 12.**
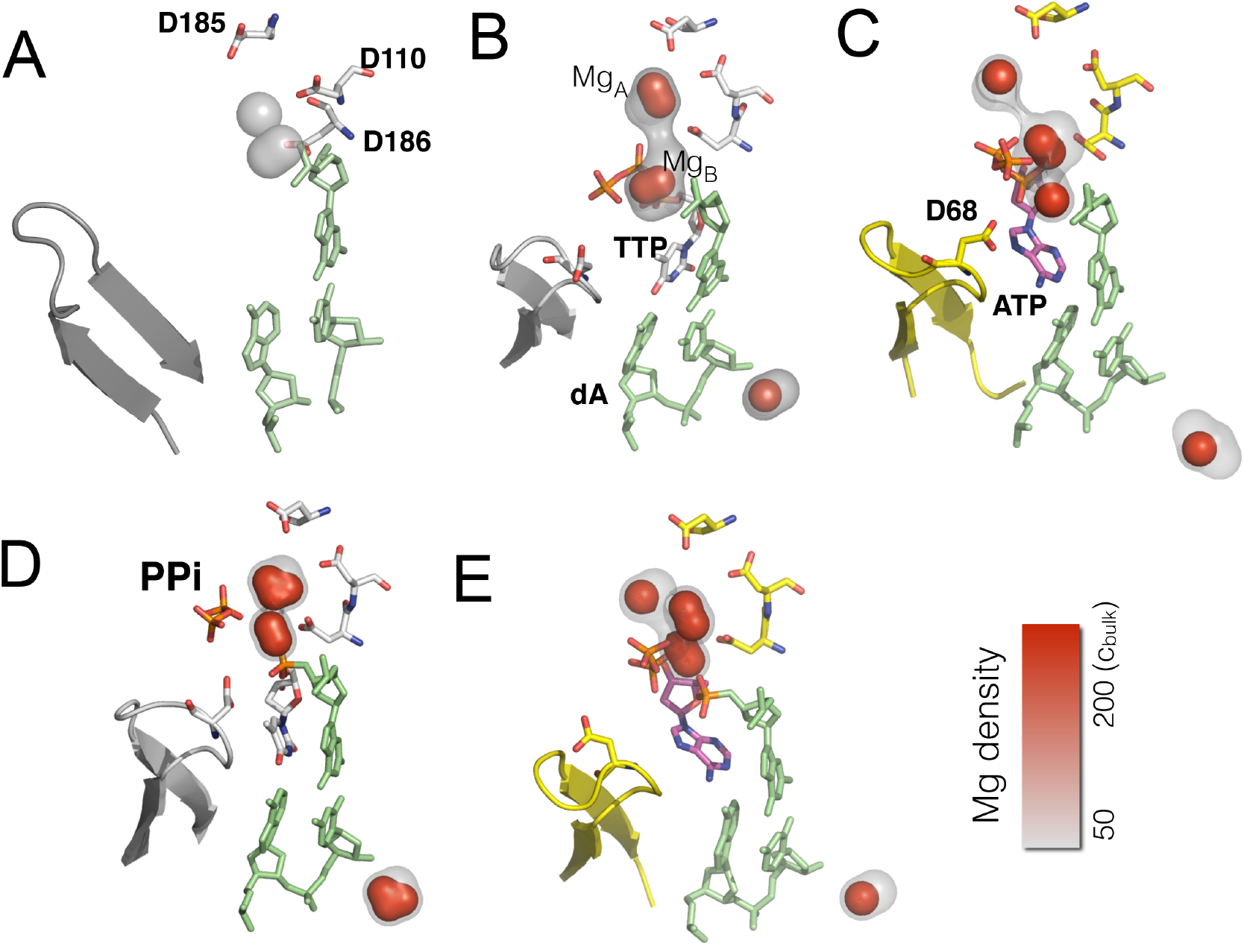
Mg ion density profiles for different steps of polymerase reaction. Local ion densities are computed by dividing the space into cubic grids. Only high-density regions are shown for clarity. 3d densities are shown with surface representation, gray to red color represents regions with ion density from *c* = 50 × *c*_bulk_ to *c* = 200 × *c*_bulk_. **A**-Corresponds to the ion density when the enzyme is in the open state, **B**-when the substrate is a matching nucleotide TTP:dA, **C**-for the mismatching nucleotide with the template ATP:dA. **D**-Shows the ion density after the chemistry step and before the PPi group dissociation from the active site for the matching nucleotide. **E-** the Mg ion density in the mismatching nucleotide.

In the open state and in the absence of Mg.dNTP, two binding sites in the vicinity of D110 and D185 are observed (Figure 12A) with local concentrations of *c* ≈ 50*c*_*bulk*_, giving rise to a 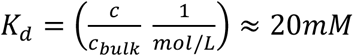. The weak Mg^2+^ binding to the open state is consistent with available crystal structures reporting the lack of observable Mg^2+^ ions in the vicinity of D110-D185 in the absence of nucleotide. This finding is also consistent with our apparent *K*_*d*_ ≈ 34 mM for nucleotide bound Mg^2+^ ion in the open state (Table 2).

The binding site between the two aspartate residues (D110 and D185) observed in the open state become more occupied when the enzyme transitions to the closed state after a correct nucleotide binds, providing a clear evidence for a dynamically changing electrostatic environment during conformational change. The equilibrium positions of the nucleotide and Mg^2+^ ion coincide well with crystal structures (Figure 9). During the simulation Mg^2+^ cations stay hydrated due to the reported (23–25) slow exchange of water from the first solvation shell of Mg^2+^. However, MgA is chelated to D110 and D185 residues in the crystal structures.

Next, we study the Mg^2+^ coordination in a mismatching ternary complex at the closed state. The Mg^2+^ binding sites in a mismatched complex (Mg.dATP with template dA) are shown in Figure 12C for comparison. Unlike the matched nucleotide, the equilibrium position of the mismatch leads to misalignment of the incoming base, consistent with our previous observations (26). Interestingly, the misalignment of the mismatch rotates the phosphate group to a bridging position between two negative charges from D68 and the terminal strand of the growing DNA. The increase in the exposed charge density in that region resulted in the formation of a *third* metal ion site due to counterion condensation. The presence of a third metal ion has been reported in repair enzymes (12,27). Our simulations suggest the possibility of a third metal ion in a high-fidelity enzyme. Care must be exercised in the interpretation of the result. Simulations suggest that the third metal ion appears only when there is a mismatch/improper alignment of the base. The dwell time of the third ion is about 8 ± 2 ns. Given a lifetime of 10 ns and diffusion-limited binding (1×10^9^ M^−1^s^−1^), we estimate a K_d_ ≅ 100 mM for the site of the third metal ion. Hence the rapid exchange of the third metal ion would make its direct observation challenging by crystallography and it is unlikely to be important under physiological conditions ([Mg^2+^] < 1 mM). Also, the weak binding affinity suggests that this site can be occupied by monovalent ions such as K^+^ or Na^+^ as they are more abundant in physiological conditions.

We also examined the Mg^2+^ ion coordination after the chemistry step, where the phosphodiester bond has just been formed and the byproduct PPi is still bound to the complex. Similar to the previous analysis we compare matched with a mismatched base pair (Figure 12D-E). The matched nucleotide and PPi created two Mg^2+^ binding sites. The binding positions are similar to the ones observed before the chemistry(Figure 12B). In contrast, a mismatched base accumulates three Mg^2+^ ions in the product complex (Figure 12E), in parallel to the mismatch before the chemistry (Figure 12C). Note that all simulations discussed in this section are independent runs started from random Mg^2+^ ion distribution. One implication of our finding is that the third metal ion provides extra electrostatic stabilization to the negatively charged PPi at the active site for the mismatch. This is consistent with our observation that PPi release is slower after the incorporation of a mismatched nucleotide (28). Further simulations are underway to study the dissociation rate of PPi group from the active site with a mismatch to complement previously published simulation studies of the release of PPi after incorporation of a correct base pair (29).

To study the kinetics and thermodynamics of the Mg^2+^ coordination to the MgA site where we measure the exchange experimentally we used the milestoning method (31). Unlike the previous cases, where spontaneous association/dissociation of Mg^2+^ ions to the negatively charged surfaces on the enzyme is directly observable in MD simulation time scale, due to relatively higher binding affinity of MgA the dissociation of magnesium ion is not within the reach of direct MD simulations. The milestoning method allowed to overcome the time-scale problem and to study the thermodynamics and kinetics of the exchange of Mg^2+^ ion from the catalytic site. Rather than following a unique Mg^2+^ ion among the pool of many in the simulation box, we monitor the *closest* Mg^2+^ ion to the unoccupied MgA binding site. This way the effect of finite concentration is taken into account as well as the indistinguishability of Mg^2+^ ions. Details of our approach is in the Methods section, here we focus on our results. The free energy change as a function of the closest Mg^2+^ ion distance is shown in Figure 13. The bound state is about 3.5 kcal/mol more stable relative to a vacant active site. The dwell time of a bound Mg^2+^ at the site on the other hand found to be 330 ns. The apparent K_d_ of 3.7 mM measured from our experiments (Figure 6A) provides an estimate of 3.2 kcal/mol. This agreement provides an important check for the reliability of our methodology. This also explains, for instance, that EDTA can instantly stop the polymerization reaction in our rapid quench experiments (32). If the Mg^2+^ dissociated more slowly, there would be a lag in stopping the reaction due to the slow Mg^2+^ dissociation.

**Figure 13.**
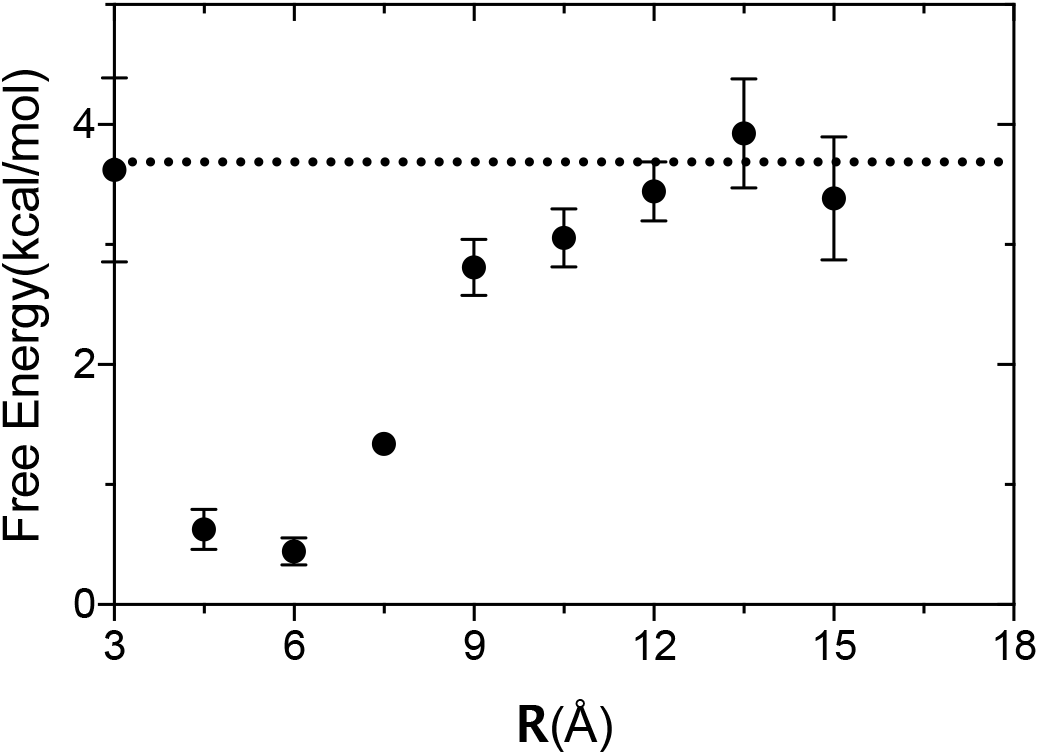
Free energy of MgA association/dissociation. Free energy profile of the closest Mg^2+^ ion to the aspartic acid pocket formed by D110 and D185. Dashed lines represent the free energy difference estimated from the experimentally measured apparent *K*_*d*_ =37 mM

## Discussion

It has been more than two decades since the general two-metal-ion mechanism for phosphoryl-transfer reactions was proposed (3). Although the metal ions could be directly observable in many crystal structures, biochemical experiments are required to establish the kinetic and thermodynamic basis for the roles of the two metal ions in specificity and catalysis. The Mg.dNTP complex is stable thermodynamically (*K*_*d*_ = 28 μM) but the metal ions exchange rapidly, complicating analysis of the roles of the second metal ion. Note that all of our experiments were conducted at concentrations of free Mg^2+^ sufficient to saturate dNTP binding. Experiments conducted at concentrations ≥0.25 mM only explore the weaker Mg^2+^ binding at the active site.

Experiments using the physiologically relevant Mg^2+^ concentrations are needed to examine the two-metal-ion mechanism based upon measurements of the kinetics of binding and catalysis. By accurate calculation of the concentrations of free Mg^2+^ and Mg.dNTP, we kinetically and thermodynamically resolved the participation of the two Mg^2+^ ions. The much weaker binding of the catalytic metal ion affords resolution of the roles of the two metal ions by titrations of activity versus free Mg^2+^. In our experiments all Mg^2+^ concentrations were above those needed to saturate the Mg.dNTP complex and provide Mg^2+^ sufficient for the counterion cloud around exposed DNA. We show kinetically that the Mg.dNTP complex binds to induce the conformational change from the open to the closed state and that the catalytic Mg^2+^ binds after the conformational change. Because the Mg-nucleotide complex is the natural substrate for many enzymes, the studies performed here also provided a more physiological general relevance about on the role of the Mg^2+^ ion *in vivo*.

It has been shown that the kinetic partitioning between the reverse conformational transition leading to release of bound nucleotide (limited by *k*_*-2*_) versus the forward reaction (*k_3_*) is a critical factor defining nucleotide specificity (*k*_*cat*_/*K*_*m*_). Our results suggested that nucleotide specificity varies as a function of the free Mg^2+^ concentration by modulating the binding of catalytic Mg^2+^ (*K*_*d,app*_ of 3.7 mM). High specificity in DNA polymerases is achieved by an induced-fit mechanism in which the enzyme closes rapidly in recognition of a correctly aligned substrate that binds tightly as it aligns catalytic residues to facilitate the binding of the catalytic Mg^2+^ to simulate the chemical reaction. A mismatched nucleotide fails to stabilize the closed state or to align catalytic residues to promote catalysis, so the mismatch is released rather than react. The catalytic Mg^2+^ contributes to high fidelity by affecting the kinetic partitioning between going forward for the chemistry versus the reverse reaction to release the substrate.

Our results resolve the controversy over the variable occupancy of the catalytic metal ion site (33), and support the theory that the reaction is catalyzed by the two-metal-ion mechanism (34). Because the second metal ion binds so weakly, it is not always observed in crystal structures at limited concentrations of Mg^2+^. The results in our study showed that Mg.dTTP binds tightly to HIV-RT in the closed state (*K*_*d,net*_ = 0.5 μM), while the catalytic Mg^2+^ binds to HIV-RT relatively weakly (*K*_*d,app*_ is 3.7 mM). The binding of Mg^2+^ to nucleotide (28 μM) is also much tighter (130-fold) than the binding of Mg^2+^ to the active site of the closed E.DNA.MgdNTP complex. We propose that the weak binding of the catalytic Mg^2+^ is an important component contributing to high fidelity. Our conclusion that two Mg^2+^ ions bind with different affinities and in sequential order to the active site of HIV-RT during the catalytic cycle is supported by available structural data. Our molecular simulations also confirm the existence of these two binding sites. Their relative binding affinities computed from simulations is consistent with our kinetic data both in the open and closed states. High-resolution x-ray crystal structures of HIV-RT have been obtained at various stages of the reaction with enzyme in complex with nucleotide analogue inhibitors (35). These structures show that HIV-RT can bind up to four functionally relevant Mg^2+^ ions, two at the HIV-RT polymerase domain and two at the RNase H domain (33,36). The interests of this study are the two Mg^2+^ in the polymerase active site exhibit significantly different affinities. There is no evidence to support the participation of a third metal ion in cognate nucleotide incorporation. However, simulations explored a weak third metal ion binding pocket in mismatch incorporation when the enzyme is in pure MgCl_2_ solution. We surmise monovalent cations abundant in physiological conditions occupy this site.

The binding of the second Mg^2+^ is required for catalysis as shown directly by our measurements, supporting the proposals first put forth in postulates of the two-metal ion mechanism (3). In addition, the second Mg^2+^ stabilizes the closed state by reducing the rate at which the enzyme opens to release the Mg.dNTP. The relatively low affinity of the second Mg^2+^ relative to the physiological concentration may provide an important contribution toward fidelity. A higher Mg^2+^ binding affinity might otherwise stabilize the binding and lead to the incorporation of a mismatched nucleotide. We are led to a model in which fidelity is largely determined by nucleotide binding to the open enzyme state and the conformational change to align the substrate at the active site, following by weak and presumably brief binding of the Mg^2+^ to stimulate catalysis. Nucleotide selectivity is based on partitioning of the closed state to go forward rather than reverse to release the bound Mg.dNTP, and the binding of the catalytic Mg^2+^ to an aligned correct substrate increases the rate of the chemical step to drive the kinetic partitioning forward.

The kinetics of the catalytic Mg^2+^ binding is supported by several experiments. The temperature dependent stopped flow experiment was repeated with three free Mg^2+^ concentrations (0.25 mM, 1 mM and 10 mM) showing no obvious effect of free Mg^2+^ on the rate of forward conformational change (*k_2_*). In addition, the global fitting of four experiments also showed that free Mg^2+^ concentration has a minimal effect on the ground state binding (*k_1_*) with an apparent *K*_*d*_≈ 34 mM for Mg binding as a competing ligand. These results demonstrate that catalytic Mg^2+^ binds after the enzyme closes. To further test this hypothesis, double mixing experiment was also performed in which Mg.dTTP was first mixed with the enzyme-DNA complex in the presence of very low free Mg^2+^ to allow the enzyme closing (Figure 7D). After a large excess of free Mg^2+^ was added at the second mixing, the chemistry occurred immediately. These results further demonstrate that the catalytic Mg^2+^ is not required for the nucleotide-induced forward conformational change but is required for catalysis.

Another approach that has been used toward dissecting the roles of the two metal ions is based on the use of the exchange-inert Rh-dNTP complex that can be purified and then mixed with the enzyme to examine the kinetics of the conformational change in the absence of Mg^2+^ (37,38). But these studies were performed before accurate measurements of the rates of the conformational change and they rely on the assumption that Rh.dNTP accurately mimics Mg.dNTP. Therefore, it was necessary to make direct measurement of the effect of free Mg^2+^ concentration on each step in the pathway, including the nucleotide-induced conformational change. As part of this study we show that the Rh.dNTP complex induces a change in structure of the enzyme from the open to the closed state in the absence of excess Mg^2+^, but with altered kinetics. The results from all three experiments are also consistent with the result from [Rh-dTTP]^2-^ binding experiment, suggesting that nucleotide bound Mg^2+^ is sufficient for inducing the enzyme closing.

Recently, it has been reported that the third Mg^2+^ is transiently bound during nucleotide incorporation, and the existence of the third Mg^2+^ was proposed to affect PPi release. The rates of the PPi release were not accurately defined in our experiments other than to show that PPi release is coincident with the observed rate of polymerization for correct nucleotide incorporation at the free Mg^2+^ concentrations ranging from 0.25 mM to 10 mM. Our previous studies on the mismatched incorporation using RNA/DNA duplex indicated the rate of PPi release is very slow (~0.03 s^−1^) and rate-limiting (39). Further studies on the mismatched incorporation using DNA/DNA duplex are needed to directly compare the rates of PPi release for correct nucleotide incorporation versus mismatched incorporation. If the rate of the PPi release is indeed very slow in mismatched nucleotide incorporation, it would suggest that nucleotide bound Mg^2+^ itself is not sufficient for facilitating the PPi release, and the proper alignment in the active site or possibly the third Mg^2+^ is required to facilitate PPi release. Pyrophosphorolysis cannot be detected with a mismatched primer/template complex over the time scale of 4 hours (Gong and Johnson, unpublished). These data suggest that the binding of Mg^2+^ and PPi do not provide sufficient energy to overcome the misalignment of the mismatched primer terminus.

Finally, we investigated nucleotide specificity (*k*_*cat*_/*K*_*m*_) at different Mg^2+^ concentrations. Our results show that nucleotide incorporation by HIVRT is highly Mg^2+^ dependent. The *k*_*cat*_/*K*_*m*_ value for dTTP incorporation decreased approximately 12-fold as the free Mg^2+^ concentration was decreased from 10 mM to 0.25 mM (Table 2). It is known that the physiological Mg^2+^ concentration varies in different cell types (40,41). For example, it is reported that the physiological concentration of free Mg^2+^ in human T lymphocytes is around 0.25 mM (42,43), but around 0.6 mM in mammalian muscle cell (44). Given the weak Mg^2+^ binding affinity, the Mg^2+^ dependent nucleotide incorporation found in our experiment may be one of the mechanisms that the host used to regulate the activities of some enzymes. On the other hand, it is possible that HIV perturbs the intracellular Mg^2+^ concentration to optimize viral replication. We have not yet measured the Mg^2+^ concentration dependence of misincorporation, which is needed to fully assess the role of Mg^2+^ concentration on fidelity.

In conclusion, we have investigated the role of each Mg^2+^ ion in the two-metal-ion mechanism by studying their binding affinities, binding mode (sequential binding or simultaneous binding), and the effects of their binding on each individual steps of the nucleotide incorporation. The studies we have performed here provided insight and detailed information about the general two-metal-ions mechanism.

## Experimental Procedures

### Mutagenesis, Expression and Purification of MDCC-labeled HIVRT

HIVRT protein was expressed, purified and labeled without resorting to the use of tagged protein as previously described (15). Briefly, cysteine mutants of the p51 (C280S) and p66 (E36C/C280S) subunits of HIVRT were separately expressed and then cells were combined to yield a 1:1 ratio of the two subunits, lysed, sonicated and then the heterodimer was purified. The protein was first purified by using the tandem Q-Sepharose and Bio-Rex70 columns, and further purified by using single-stranded DNA (ssDNA) affinity column. The protein was then labeled with the MDCC (7-Diethylamino-3-[*N*-(2-maleimidoethyl) carbamoyl]coumarin) from Sigma-Aldrich. Unreacted MDCC was removed by ion exchange using a Bio-Rex70 column. After purification, ‘Coomassie Plus’ protein assay was used to estimate the purified protein concentration (Thermofisher). In addition, an active site titration was performed to determine the active site concentrations of the purified protein (14), which was used in all subsequent experiments.

### Preparation of DNA Substrates for Kinetic Studies

The 25/45nt and 25ddA/45nt DNA substrates were purchased from Integrated DNA Technologies, using the following sequences:

25nt: 5’-GCCTCGCAGCCGTCCAACCAACTCA−3’
45nt: 5’-GGACGGCATTGGATCGACGATGAGTTGGTTGGACGGCTGCGAGGC −3’

The oligonucleotides were annealed by heating at 95°C for 5 minutes, followed by slow cooling to room temperature. For making radiolabeled primer, the 25-nt oligonucleotide was labeled at the 5’ end by γ-^32^P ATP (PerkinElmer) using T4 polynucleotide kinase (NEB).

### Quench Flow Kinetic Assays

Rapid chemical-quench-flow experiments were performed by mixing a preformed enzyme-DNA complex (using radio-labeled DNA primer) with various concentrations of incoming nucleotide using a KinTek RQF-3 instrument (KinTek Corp). The reaction was quenched by the addition of 0.5 M EDTA at varying time points. Products were collected and separated on 15% denaturing PAGE (acrylamide (1:19 bisacrylamide), 7M Urea). Results were then analyzed using ImageQuant 6.0 software (Molecular Dynamics).

### Stopped Flow Kinetic Assays

The stopped-flow measurements were performed by rapidly mixing an enzyme-DNA complex (using MDCC-labeled HIVRT) with various concentrations of incoming nucleotide. The time dependence of fluorescence change upon nucleotide binding and incorporation was monitored using an AutoSF-120 stopped-flow instrument (KinTek Corp, Austin, TX) by exciting the fluorophore at 425 nm and monitoring the fluorescence change at 475 nm using a band-pass filter with a 25 nm bandwidth (Semrock).

### Global Fitting of Multiple Experiments

The kinetic parameters governing each step leading to nucleotide incorporation were obtained by globally fitting four experiments (stopped-flow, chemical-quench, nucleotide off-rate, and PPi release) using the model shown in Scheme 1 with *KinTek Explorer* software (KinTek Corp, Austin, TX). Fitspace confidence contour analysis was also performed to estimate standard errors (45,46).

### Free Mg^2+^ Concentration Calculation

To calculate the free Mg^2+^ concentration in solution, EDTA (500 μM) was used as a buffering system. The main equilibria that affect free Mg^2+^ concentration are the equilibrium constants for Mg^2+^ binding to dNTP, Mg^2+^ binding to EDTA, and the equilibrium for protonation of dNTP (shown in Table 7). Calculation of the free Mg^2+^ concentration from starting total concentrations of Mg^2+^, dNTP and EDTA and the pH requires an iterative approach to solve simultaneously the equilibria involved. However, we simplified the problem by specifying the desired free Mg^2+^ concentration and total EDTA concentration, and then calculating the total concentrations of Mg^2+^ and dNTP that must be added to the solution using a simplified set of equations:

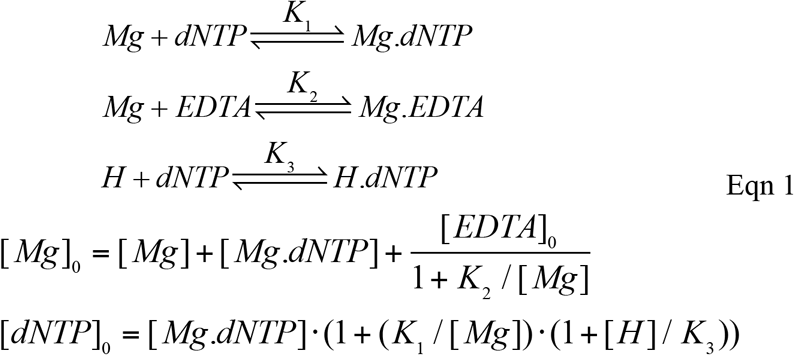

**Table 7.** Equilibrium constants used for the calculation of concentrations of free magnesium and Mg.dNTP concentrations. The association and dissociation constants, *K*_*a*_ and *K*_*d*_, respectively, were obtained from (Storer AC, *et al.,* 1976; Martell *et al.,* 1964) (4,5).

| Equilibrium | $K_a$ ($M^{-1}$) | $K_d$ ($\mu M$) |
| --- | --- | --- |
| $Mg^{2+} + ATP^{4-} \rightleftharpoons Mg.ATP^{2-}$ | 34800 | 28.7 |
| $H^+ + ATP^{4-} \rightleftharpoons H.ATP^{3-}$ | $1.09 \times 10^7$ | $9.17 \times 10^{-2}$ |
| $H^+ + H.ATP^{3-} \rightleftharpoons H_2.ATP^{2-}$ | 8500 | 118 |
| $Mg^{2+} + H.ATP^{3-} \rightleftharpoons Mg.H.ATP^{-}$ | 542 | 1845 |
| $Mg^{2+} + EDTA^{4-} \rightleftharpoons Mg.EDTA^{2-}$ | $4 \times 10^8$ | $0.25 \times 10^{-2}$ |
| $H^+ + EDTA^{4-} \rightleftharpoons H.EDTA^{3-}$ | $1.66 \times 10^{10}$ | $6 \times 10^{-5}$ |
| $H^+ + H.EDTA^{3-} \rightleftharpoons H_2.EDTA^{2-}$ | $1.58 \times 10^6$ | 0.633 |

To confirm the accuracy of our approximation, the reactions were simulated with all equilibria (Table 7) (4,5) using *KinTek Explorer* software. In this case, starting total concentrations of Mg^2+^, dNTP and EDTA were entered, and the free Mg^2+^ concentration was directly calculated after the system reached equilibrium. This kinetic approach to reach equilibrium circumvents the typical semi-random search for a mathematical solution and yet still affords a simultaneous solution of the multiple equilibria.

### Calculation of steady state kinetic parameters

Steady state kinetic parameters were calculated from the intrinsic rate constants using the following equations according to Scheme 1, simplified by the known fast product release (*k*_*4*_ ⨠ *k_3_*). The initial ground-state binding was modeled as a rapid equilibrium with *k_1_* = 100 μM^−1^s^−1^. Estimates of the remaining rate constants were then used to calculate the steady state kinetic parameters.

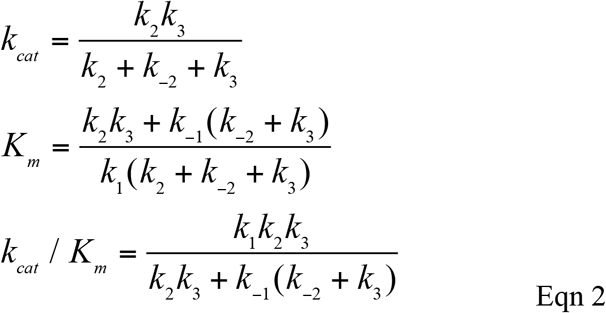

#### Initial state of MD simulation models

Initial states of the ED-Mg.dNTP ternary complex were based on the open (1j5o) and closed state(1rtd) structures of HIV1-RT from the protein databank (47). Five independent simulation setups were prepared as follows: I) open state enzyme with no bound dNTP, II) closed state enzyme with matching nucleotide, [Mg.dTTP]^−2^ opposite to a templating base Adenine forming dTTP:dA pair, III) closed state enzyme with a mismatching nucleotide, [Mg.ATP]^−2^ opposite to templating DNA forming dATP:dA pair, IV) closed state enzyme after the chemistry step where matching nucleotide is added to the DNA strand and pyrophosphate (PPi) group is bound, V) closed state enzyme after chemistry step where mismatching nucleotide added and PPi is bound.

#### MD simulation set up

Molecular dynamics simulations were performed using the GROMACS suit of programs (48). Each set up was first energy minimized with steepest descent method for 5000 steps followed by solvation with explicit water with a minimum of 12 Å thickness from the surface of the complex, giving rise to a simulation box of about 126.6 × 125.2 × 112.7 Å^3^. To neutralize the simulation box and to mimic experimental conditions we added 51 Mg^2+^ and 63 Cl^−^ by randomly replacing water molecules with ions, resulting in 30∓1mM free Mg^2+^ in the bulk after equilibriation. Water was represented by SPC/E model(49). Protein, DNA, and Cl^−^ molecules were represented by default Amber03 forcefield parameters (50). For Mg^2+^ ions we used recently developed parameter (25) that shows better agreement with solution exchange rates. The parameters of dTTP, dATP, and PPi were adopted from Amber forcefield while charges were computed from quantum mechanics as explained in our earlier work (26).

We compute the long-range interactions with a distance cutoff of 12Å with dispersion correction for van der Waals interactions (51) and Particle Mesh Ewald summation method (52) for electrostatic. Electrostatic interactions were computed with a grid spacing of 1.15, 1.16, and 1.12 Å in directions *x*, *y*, and *z*. Equations of motion were integrated by Leap-frog integrator (53) with a time step of 2fs. All bonds were constrained using LINCS (54) algorithm.

Following the energy minimization for each solvated system we conducted a two-step equilibration process: the first equilibration involves finding the volume of the box that give rise to 1 atm pressure at 310K, the second equilibration allowed water and ions to equilibrate around the complex. In detail, we sampled the conformations for 2 ns from Isothermal Isobaric ensemble (NPT) using Parrinello–Rahman scheme (55) and the temperature was kept constant by velocity scaling (52). The positions of heavy atoms of the solute were restrained using harmonic potential with a stiffness constant of 1000 kJ/nm^2^. Using the last frame of the simulation as starting point, we employed 200 ns long constant volume and temperature (NVT) simulation for solvent equilibration. In this stage, stiffness constant of the position restraints was decreased to 50 J/nm^2^ to allow local adjustments in the enzyme-substrate complex. The last frame of the trajectory was used as a starting point for unconstrained MD simulations where we compute our observables. For sampling equilibrium configurations from NVT ensemble we employed a minimum of 300 ns long simulation in each setup described above. We removed the translational and rotational degrees of the enzyme for every 10 ps and coordinates of atom positions were recorded for each picosecond for data analysis.

#### Computing ion density

To study Mg^2+^ distribution around the complex, we divided the simulation box into cubic grids of 1Å in each direction and we computed the average Mg^2+^ ion occupancy at each grid. From the occupancy, local ion concentrations and free Mg^2+^ concentration at the bulk were computed. To estimate bulk concentration, we averaged the concentrations of all grids that are 12 Å or more away from the enzyme surface.

#### Computing the kinetics and thermodynamics of Mg_A_

To study the kinetics and thermodynamics of Mg^2+^ coordination to catalytic site we developed a method employing Milestoning approach. In milestoning approach we partition the phase space into milestones. Using trajectory fragments, we estimate the stationary flux between milestones. Details of the milestoning approach employed can be found in Ref (56,57). From the stationary flux we compute the average mean first passage time, (MFPT), 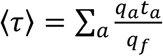, where *q*_*a*_ is the stationary flux of milestone *a*, *t*_a_ is the average dwell time at *a*, and *q*_*f*_ is the flux to the product state. In addition, the stationary fluxes can be used to extract the free energy of each milestone using *F*_a_ = −*k*_*b*_*T* ln(*q*_a_*t*_a_).

Milestoning in our study requires a reasonable pathway and a reaction coordinate that quantify the progress of the ion migration process. To obtain Mg^2+^ ion binding pathway we created 25 configurations from equilibrium MD simulations obtained from *Set up 2* described above. As in all of these configurations there was a Mg ion in the MgA site. We moved the ion to a random place in the simulation box to create a vacancy at the catalytic site. After 100 ps of solvent equilibration of the entire system we monitored the diffusion of all remote Mg^2+^ ions by computing the closest magnesium ion to the carboxylic acids atoms of the pocket created by D110 and D185. We employed about 100 ns sampling of ion conformations at various initial states, totaling of 2.5 microseconds of simulation time. These trajectories later used to compute the transition kernel, *K*_ab_ where *a* and *b* corresponds to neighboring milestone indices. The milestones are distributed between the initial end states in 1Å intervals. We ensure the convergence of the transition probabilities. To study the association rate, we assigned absorbing boundary conditions to the two Mg bound states, defined by *R* = 4Å (product) and a reflecting boundary condition to *R* = *R*_bulk_ (reactant). Here, *R*_bulk_ is defined as the average distance between two Mg^2+^ if they uniformly distributed in solution. *R*bulk is ~16 Å in 30 mM free Mg^2+^ concentration. Knowing that the dNTP bound Mg is already at the active site this distance is the average distance that one can find another Mg ion. We used association pathway that is accessible to brute force MD to compute the kinetics of the dissociation process. For dissociation rates the milestoning equations are solved with boundary conditions reversed.

## Data Availability

KinTek Explorer mechanism files used to fit data and refined HIV RT structure files used for MD simulations are available upon request.

## Acknowledgement

Supported by grants from the Welch Foundation (F-1604 to K.A.J.), The National Institutes of Health (R01GM114223 and R01AI110577 to K.A.J.; and R01GM104896 and R01GM125882 to Y.J.Z), and AD181 faculty research grant (to S.K.). We wish to thank for Dr. William H. Konigsberg, who kindly provided [Rh-dTTP]^2^. MD simulations were carried out on the High-Performance Computing resources at New York University Abu Dhabi.

## Abbreviations

HIVRT: Human Immunodeficiency Virus Reverse Transcriptase
dTTP: Thymidine triphosphate
dNTP: deoxynucleoside triphosphate
Mg_B_: nucleotide-bound Mg^2+^
Mg_A_: Catalytic Mg^2+^
ED: enzyme-DNA complex
ED_dd_: enzyme-DNA complex with a dideoxy-terminated primer strand
MDCC: 7-diethylamino-3-((((2-maleimidyl)ethyl)amino)carbonyl) coumarin.

## References

1. Yang, L. J., Arora, K., Beard, W. A., Wilson, S. H., and Schlick, T. (2004) Critical role of magnesium ions in DNA polymerase beta’s closing and active site assembly. J Am Chem Soc 126, 8441–8453

2. Fenstermacher, K. J., and DeStefano, J. J. (2011) Mechanism of HIV reverse transcriptase inhibition by zinc: formation of a highly stable enzyme-(primer-template) complex with profoundly diminished catalytic activity. The Journal of biological chemistry 286, 40433–40442

3. Steitz, T. A., and Steitz, J. A. (1993) A general two-metal-ion mechanism for catalytic RNA. Proc Natl Acad Sci U S A 90, 6498–6502

4. Martell, L. (1964) Stability constants of metal-ion complex. The Chemical Society

5. Storer, A. C., and Cornish-Bowden, A. (1976) Concentration of MgATP2-and other ions in solution. Calculation of the true concentrations of species present in mixtures of associating ions. Biochem J 159, 1–5

6. Freudenthal, B. D., Beard, W. A., Shock, D. D., and Wilson, S. H. (2013) Observing a DNA polymerase choose right from wrong. Cell 154, 157–168

7. Atis, M., Johnson, K. A., and Elber, R. (2017) Pyrophosphate Release in the Protein HIV Reverse Transcriptase. The journal of physical chemistry. B 121, 9557–9565

8. Steitz, T. A. (1999) DNA polymerases: structural diversity and common mechanisms. The Journal of biological chemistry 274, 17395–17398

9. Nakamura, T., Zhao, Y., Yamagata, Y., Hua, Y. J., and Yang, W. (2012) Watching DNA polymerase eta make a phosphodiester bond. Nature 487, 196–201

10. Yang, W., Weng, P. J., and Gao, Y. (2016) A new paradigm of DNA synthesis: three-metal-ion catalysis. Cell Biosci 6, 51

11. Stevens, D. R., and Hammes-Schiffer, S. (2018) Exploring the Role of the Third Active Site Metal Ion in DNA Polymerase eta with QM/MM Free Energy Simulations. J Am Chem Soc 140, 8965–8969

12. Tsai, M. D. (2019) Catalytic mechanism of DNA polymerases-Two metal ions or three? Protein Sci 28, 288–291

13. Kellinger, M. W., and Johnson, K. A. (2011) Role of induced fit in limiting discrimination against AZT by HIV reverse transcriptase. Biochemistry 50, 5008–5015

14. Tsai, Y. C., and Johnson, K. A. (2006) A new paradigm for DNA polymerase specificity. Biochemistry. 45, 9675–9687

15. Kellinger, M. W., and Johnson, K. A. (2010) Nucleotide-dependent conformational change governs specificity and analog discrimination by HIV reverse transcriptase. Proc Natl Acad Sci U S A 107, 7734–7739

16. Johnson, K. A. (2019) Kinetic Analysis for the New Enzymology: Using computer simulation to learn kinetics and solve mechanisms., KinTek Corporation, Austin, USA

17. Huang, H. F., Chopra, R., Verdine, G. L., and Harrison, S. C. (1998) Structure of a covalently trapped catalytic complex of HIV-I reverse transcriptase: Implications for drug resistance. Science 282, 1669–1675

18. Hanes, J. W., and Johnson, K. A. (2008) Real-time measurement of pyrophosphate release kinetics. Anal Biochem 372, 125–127

19. Hanes, J. W., and Johnson, K. A. (2007) A novel mechanism of selectivity against AZT by the human mitochondrial DNA polymerase. Nucleic acids research 35, 6973–6983

20. Brune, M., Hunter, J. L., Howell, S. A., Martin, S. R., Hazlett, T. L., Corrie, J. E., and Webb, M. R. (1998) Mechanism of inorganic phosphate interaction with phosphate binding protein from Escherichia coli. Biochemistry 37, 10370–10380

21. Manning, G. S. (1984) Limiting Laws and Counterion Condensation in Poly-electrolyte Solutions 8. Mixtures of Counterions, Specific Selectivity, and Valence Selectivity. J Phys Chem-Us 88, 6654–6661

22. Kirmizialtin, S., Silalahi, A. R. J., Elber, R., and Fenley, M. O. (2012) The Ionic Atmosphere around A-RNA: Poisson-Boltzmann and Molecular Dynamics Simulations. Biophys J 102, 829–838

23. Bleuzen, A., Pittet, P. A., Helm, L., and Merbach, A. E. (1997) Water exchange on magnesium(II) in aqueous solution: a variable temperature and pressure O-17 NMR study. Magn Reson Chem 35, 765–773

24. Lee, Y., Thirumalai, D., and Hyeon, C. (2017) Ultrasensitivity of Water Exchange Kinetics to the Size of Metal Ion. J Am Chem Soc 139, 12334–12337

25. Allner, O., Nilsson, L., and Villa, A. (2012) Magnesium Ion-Water Coordination and Exchange in Biomolecular Simulations. J Chem Theory Comput 8, 1493–1502

26. Kirmizialtin, S., Nguyen, V., Johnson, K. A., and Elber, R. (2012) How Conformational Dynamics of DNA Polymerase Select Correct Substrates: Experiments and Simulations. Structure 20, 618–627

27. Yang, W., Weng, P. J., and Gao, Y. (2016) A new paradigm of DNA synthesis: three-metal-ion catalysis. Cell Biosci 6

28. Li, A., Gong, S. Z., and Johnson, K. A. (2016) Rate-limiting Pyrophosphate Release by HIV Reverse Transcriptase Improves Fidelity. Journal of Biological Chemistry 291, 26554–26565

29. Atis, M., Johnson, K. A., and Elber, R. (2017) Pyrophosphate Release in the Protein HIV Reverse Transcriptase. J Phys Chem B 121, 9557–9565

30. Rungrotmongkol, T., Mulholland, A. J., and Hannongbua, S. (2014) QM/MM simulations indicate that Asp185 is the likely catalytic base in the enzymatic reaction of HIV-1 reverse transcriptase. Medchemcomm 5, 593–596

31. Faradjian, A. K., and Elber, R. (2004) Computing time scales from reaction coordinates by milestoning. J Chem Phys 120, 10880–10889

32. Patel, S. S., Wong, I., and Johnson, K. A. (1991) Pre-Steady-State Kinetic-Analysis of Processive Dna-Replication Including Complete Characterization of An Exonuclease-Deficient Mutant. Biochemistry 30, 511–525

33. Cowan, J. A., Ohyama, T., Howard, K., Rausch, J. W., Cowan, S. M., and Le Grice, S. F. (2000) Metal-ion stoichiometry of the HIV-1 RT ribonuclease H domain: evidence for two mutually exclusive sites leads to new mechanistic insights on metal-mediated hydrolysis in nucleic acid biochemistry. J Biol Inorg Chem 5, 67–74

34. Klumpp, K., Hang, J. Q., Rajendran, S., Yang, Y., Derosier, A., Wong Kai In, P., Overton, H., Parkes, K. E., Cammack, N., and Martin, J. A. (2003) Two-metal ion mechanism of RNA cleavage by HIV RNase H and mechanism-based design of selective HIV RNase H inhibitors. Nucleic acids research 31, 6852–6859

35. Sarafianos, S. G., Clark, A. D., Jr., Das, K., Tuske, S., Birktoft, J. J., Ilankumaran, P., Ramesha, A. R., Sayer, J. M., Jerina, D. M., Boyer, P. L., Hughes, S. H., and Arnold, E. (2002) Structures of HIV-1 reverse transcriptase with pre-and post-translocation AZTMP-terminated DNA. EMBO J. 21, 6614–6624

36. Cristofaro, J. V., Rausch, J. W., Le Grice, S. F., and DeStefano, J. J. (2002) Mutations in the ribonuclease H active site of HIV-RT reveal a role for this site in stabilizing enzyme-primer-template binding. Biochemistry 41, 10968–10975

37. Bakhtina, M., Lee, S., Wang, Y., Dunlap, C., Lamarche, B., and Tsai, M. D. (2005) Use of viscogens, dNTPalphaS, and rhodium(III) as probes in stopped-flow experiments to obtain new evidence for the mechanism of catalysis by DNA polymerase beta. Biochemistry 44, 5177–5187

38. Lee, H. R., Wang, M., and Konigsberg, W. (2009) The reopening rate of the fingers domain is a determinant of base selectivity for RB69 DNA polymerase. Biochemistry 48, 2087–2098

39. Li, A., Gong, S., and Johnson, K. A. (2016) Rate-limiting Pyrophosphate Release by HIV Reverse Transcriptase Improves Fidelity. The Journal of biological chemistry 291, 26554–26565

40. Valberg, L. S., Holt, J. M., Paulson, E., and Szivek, J. (1965) Spectrochemical Analysis of Sodium, Potassium, Calcium, Magnesium, Copper, and Zinc in Normal Human Erythrocytes. J Clin Invest 44, 379–389

41. Walser, M. (1967) Magnesium metabolism. Ergeb Physiol 59, 185–296

42. Delva, P., Pastori, C., Degan, M., Montesi, G., and Lechi, A. (1998) Intralymphocyte free magnesium and plasma triglycerides. Life Sci 62, 2231–2240

43. Delva, P., Pastori, C., Degan, M., Montesi, G., and Lechi, A. (2004) Catecholamine-induced regulation in vitro and ex vivo of intralymphocyte ionized magnesium. J Membr Biol 199, 163–171

44. Tashiro, M., and Konishi, M. (1997) Basal intracellular free Mg^2+^ concentration in smooth muscle cells of guinea pig tenia cecum: intracellular calibration of the fluorescent indicator furaptra. Biophys J 73, 3358–3370

45. Johnson, K. A., Simpson, Z. B., and Blom, T. (2009) FitSpace Explorer: An algorithm to evaluate multidimensional parameter space in fitting kinetic data. Anal Biochem 387, 30–41

46. Johnson, K. A., Simpson, Z. B., and Blom, T. (2009) Global Kinetic Explorer: A new computer program for dynamic simulation and fitting of kinetic data. Anal Biochem 387, 20–29

47. Berman, H. M., Westbrook, J., Feng, Z., Gilliland, G., Bhat, T. N., Weissig, H., Shindyalov, N., and Bourne, P. E. (2000) The Protein Data Bank. Nucleic Acids Res 28, 235–242

48. Mark James Abraham, T. M., Roland Schulz, Szilárd Páll, Jeremy C. Smith, Berk Hess, Erik Lindahl,. (2015) GROMACS: High performance molecular simulations through multi-level parallelism from laptops to supercomputers. SoftwareX 1-2, 19–25

49. Berendsen, H. J. C., Grigera, J. R., and Straatsma, T. P. (1987) THE MISSING TERM IN EFFECTIVE PAIR POTENTIALS. Journal of Physical Chemistry 91, 6269–6271

50. Duan, Y., Wu, C., Chowdhury, S., Lee, M. C., Xiong, G. M., Zhang, W., Yang, R., Cieplak, P., Luo, R., Lee, T., Caldwell, J., Wang, J. M., and Kollman, P. (2003) A point-charge force field for molecular mechanics simulations of proteins based on condensed-phase quantum mechanical calculations. J Comput Chem 24, 1999–2012

51. Wennberg, C. L., Murtola, T., Hess, B., and Lindahl, E. (2013) Lennard-Jones Lattice Summation in Bilayer Simulations Has Critical Effects on Surface Tension and Lipid Properties. J. Chem. Theory Comput 9, 3527–3537

52. G. Bussi, D. D. a. M. P. (2007) Canonical sampling through velocity-rescaling. J Chem Phys 126

53. Van Gunsteren, W. F., and Berendsen, H. J. C. (1988) A Leap-Frog Algorithm for Stochastic Dynamics. Mol Simulat 1, 173–185

54. Hess, B. (2008) P-LINCS: A parallel linear constraint solver for molecular simulation. J Chem Theory Comput 4, 116–122

55. Parrinello, M. R., A. Polymorphic transitions in single crystals: A new molecular dynamics method. Journal of Applied Physics 52, 7182–7190

56. Kirmizialtin, S., and Elber, R. (2011) Revisiting and Computing Reaction Coordinates with Directional Milestoning. J Phys Chem A 115, 6137–6148

57. West, A. M. A., Elber, R., and Shalloway, D. (2007) Extending molecular dynamics time scales with milestoning: Example of complex kinetics in a solvated peptide. J Chem Phys 126

